# A mathematical model of viral oncology as an immuno-oncology instigator

**DOI:** 10.1101/429233

**Authors:** Tyler Cassidy, Antony R. Humphries

## Abstract

We develop and analyse a mathematical model of tumour-immune interaction that explicitly incorporates heterogeneity in tumour cell cycle duration by using a distributed delay differential equation. Our necessary and sufficient conditions for local stability of the cancer free equilibrium completely characterise the importance of tumour-immune interaction in disease progression. Consistent with the immunoediting hypothesis, we show that decreasing tumour-immune interaction leads to tumour expansion. Finally, we show that immune involvement is crucial in determining the long-term response to viral therapy.

## 1 Introduction

Malignant tumours contain a highly heterogeneous population of cells that have distinct genotypes and reproductive abilities [Bell and McFadden, 2014; Lichty et al., 2014]. The heterogeneous nature of tumours is mirrored in the reproduction speed of malignant cells. Most existing mathematical models greatly simplify the impact of heterogeneity in cell cycle times by either neglecting the cell cycle or assuming that all tumour cells have identical cell cycle durations. We will account for the range of cell cycle durations by deriving a mathematical model of tumour growth using a delay differential equation (DDE) with a distribution of delays. This is, to our knowledge, a novel method of considering the heterogeneity present in malignant tumours and presents a physiologically realistic model of tumour expansion.

Distributed DDEs model a continuum of cell cycle durations that belong to an interval of physiologically realistic values, with durations distributed according to a probability density function (PDF). Representing the time length of the cell cycle by a distributed DDE explicitly allows for variability in cell cycle duration. This contrasts with discrete DDEs, where the discrete delay represents the cell cycle duration which is taken to be the same for all tumour cells. Thus discrete delays implicitly assume homogeneity of the tumour cell cycle duration which limits the physiological relevance of such models.

The human immune system attempts to eradicate malignant cells and inhibit tumour establishment [Hallam et al., 2009; Hoos et al., 2011]. We study this phenomenon by explicitly including tumour-immune interaction in our mathematical model. Analysis of this model shows that there is a threshold tumour size below which the immune system successfully prevents tumour establishment and quantifies the role of immune surveillance in tumour establishment and growth.

Therapeutic strategies under development attempt to exploit the immune system to eradicate malignant tumours via immuno-oncology and genetically engineered oncolytic viruses [Cassady et al., 2016; Chiocca and Rabkin, 2015; Hoos et al., 2011; Lawler and Chiocca, 2015]. Oncolytic viruses are designed to exploit the high reproductive rate characteristic of malignant tumours and preferentially infect cancerous cells. Immune regulated death of infected tumour cells releases tumour specific antigens that signal the immune system [Breitbach et al., 2016]. We incorporate oncolytic viral therapy into our mathematical model to study how these viruses can prime the immune system to eliminate tumours.

The release of tumour specific antigens induces a long-lasting immune response that causes tumour regression which persists after resolution of the infection [Bourgeois-Daigneault et al., 2016]. Consequently, oncolytic viruses have recently been recast as instigators of immuno-oncology and are being engineered to induce immune recruitment. For example, in 2015, the United States Food and Drug Administration approved a modified herpes virus that promotes granulocyte-macrophage colony-stimulating factor production and resulting anti-tumour immunity for treatment of melanoma [Bommareddy et al., 2017].

Mathematical models have been used extensively to understand and predict tumour growth and tumour-immune interactions (see Santiago et al. [2017]; Walker and Enderling [2016]; Wodarz [2016] for reviews). Existing models range from formulations as ordinary differential equations (ODEs) [Idema et al., 2010; Kim et al., 2015; Kirschner and Panetta, 1998; MacNamara and Eftimie, 2015], to partial differential equations [Hillen et al., 2013; Malinzi et al., 2017] and discrete DDEs [Liu et al., 2007; Mahasa et al., 2017; Villasana and Radunskaya, 2003].

Crivelli et al. [2012] developed and analysed a discrete DDE model of tumour growth and viral oncology. The Crivelli model is simple enough to be analytically tractable while retaining important physiological aspects of tumour growth and oncolytic viral therapy, but neglects the role of the immune system in tumour eradication. Crivelli et al. [2012] model the interaction of virions and tumour cells by using a non-differentiable function which significantly complicates the analysis of the model. This contact function allows for viral therapy alone to drive tumour remission in their model, without interaction with the immune system.

We develop a tumour growth and viral oncology model which incorporates immune recruitment to drive tumour clearance. Our model is partly based on the Crivelli model but augments and generalises it in very significant ways. We explicitly model phagocytosis of the tumour cells, and cytokine driven phagocyte recruitment. As mentioned, we also include a distribution of cell cycles times for the tumour cells which results in a DDE with distributed delays. The inclusion of a heterogeneous cell cycle duration is more realistic than models with a discrete delay, because a discrete delay is equivalent to assuming that that every cell in the tumour has a constant and identical cell cycle duration. We show the explicit link between our work and Crivelli et al. [2012] in Appendix A.

The distributed DDE tumour-immune model is developed in full generality in Section 2. In Section 3, we prove that solutions of the initial value problem evolving from non-negative initial data remain nonnegative. Next, in Theorem 3.3, we determine a condition for treatment free extinction of the tumour that quantifies the link between immune involvement and disease progression. Our results show that immune involvement is crucial in controlling tumour growth. As a direct consequence, we show in Corollary 3.4 that homogeneous tumours are less robust than tumours with heterogeneous cell cycle durations. Finally, by showing the existence of a cancer-immune co-existence equilibrium in Theorem 3.5, we establish a direct link between the minimal viable tumour size and the immune killing capacity that is consistent with the immunoediting hypothesis of tumour progression [Mittal et al., 2014]. In Section 4, by deriving a variant of the linear chain technique, we prove that the distributed DDE is equivalent to a finite dimensional ODE. We end Section 4 by simulating viral oncology treatment and illustrating the previously derived stability results. Our simulations show the existence of a transcritical bifurcation where the unstable nonzero equilibrium acts as a separatrix between tumour extinction and growth. Biologically, this result implies that treatment strategies that force the malignant tumour across the separatrix will eradicate the tumour. Moreover, we show that sufficiently strong immune involvement can counteract aggressive tumour growth and lead to tumour extinction without treatment. Finally, we discuss our results in Section 5.

## 2 Model Development

Our model of tumour-immune interaction is given by the system of differential equations

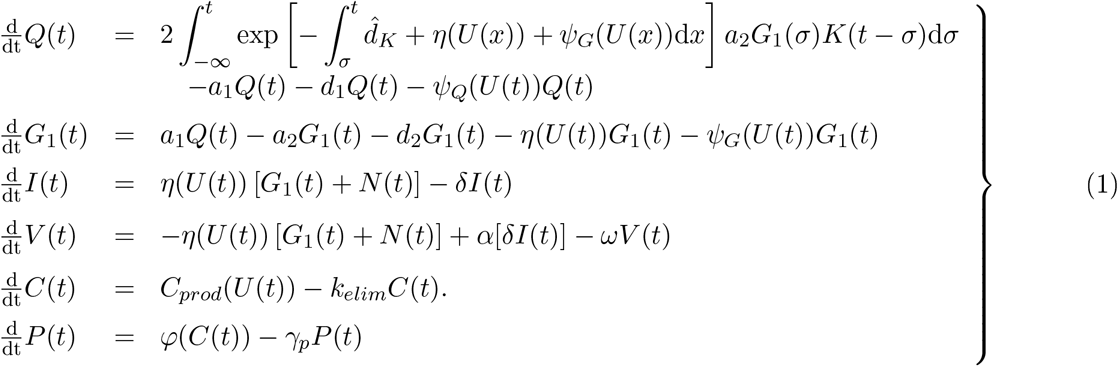

In equation (1), *Q*(*t*) and *G*_1_(*t*) denote the quiescent and proliferative phase tumour cells. The cytokine concentration is denoted by *C*(*t*), and the phagocyte concentration in the tumour microenvironment by *P*(*t*). Finally, *V*(*t*) is the concentration of oncolytic virions and *I*(*t*) is the number of infected tumour cells.

In the Burns and Tannock [1970] model of the cell cycle, *Q*(*t*) corresponds to cells in the *G*_0_ phase while *G*_1_(*t*) corresponds to the *G*_1_ phase. We consider the *S, G*_2_ and *M* phases to be the active phases of the cell cycle, which we model as a process rather than as populations.

In a similar manner to Crivelli et al. [2012]; Dawson and Hillen [2006] and Liu et al. [2007], we use a constant transition rate from *G*_0_ to *G*_1_. The *G*_1_-*S* and the *G*_2_-*M* checkpoints have been explored as targets of emerging cancer treatment [Dominguez-Brauer et al., 2015; Matheson et al., 2016; Visconti et al., 2016]. By separating the *G*_1_ phase from the *S, G*_2_ and *M* phases, our model could be easily adapted to include the precise effects of interventions that arrest the cell cycle at the *G*_1_-*S* checkpoint, such as cyclin dependent kinase inhibitors [Dominguez-Brauer et al., 2015]. It would also be possible to incorporate drug induced cell cycle arrest at the *G*_2_-*M* checkpoint by decreasing mitotic output (without effecting transition across the *G*_1_-*S* checkpoint), similar to emerging treatments discussed by Dominguez-Brauer et al. [2015]; Visconti et al. [2016].

We denote by *N*(*t*) the total number of cells in the active portion (the *S, G*_2_ and *M* phases) of the cell cycle, given by

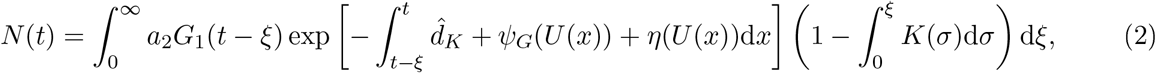
as derived in Appendix B. In equations (1) and (2) the distribution of the duration of the active phase of the cell cycle is described by the PDF *K*(*t*). We do not choose a specific distribution in our model; see Section 2.1 for a discussion of the properties of *K*(*t*).

The functions *η*(*U*(*t*)), *ψ_Q_*(*U*(*t*)), *ψ_G_*(*U*(*t*)), *φ*(*C*(*t*)), and *C_prod_*(*U*(*t*)) in equation (1) are defined in equations (8), (10), (11) and (13). To simplify notation, we denote the vector

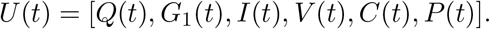

The distributed DDE is given initial data *Q*(*t*_0_), *I*(*t*_0_), *C*(*t*_0_) and [*G*_1_(*s*), *V*(*s*), *P*(*s*)] = [*ϕ_G_*(*s*), *ϕ_V_*(*s*), *ϕ_P_*(*s*)] for *s* ∈ (−∞, *t*_0_] for integrable functions *ϕ_G_*(*s*), *ϕ_V_*(*s*) *ϕ_P_*(*s*) to create an initial value problem. For simplicity, we take *t*_0_ = 0.

We derive equation (1) in three steps. First, we consider tumour growth in the absence of immune interaction and viral therapy in Section 2.1. Tumour heterogeneity is explicitly accounted for by using a distributed cell cycle time length. The tumour growth equations are derived keeping in mind the eventual use of the model to describe the impact of an RNA oncolytic virus on tumour growth. Next, in Section 2.2, we derive the tumour-immune interaction and incorporate immunosurveillance into the tumour growth model. The graphical representation of the tumour-immune growth model is given in Figure 1. Finally, by including viral therapy and immune recruitment in Section 2.3, we arrive at equation (1).

**Figure 1:**
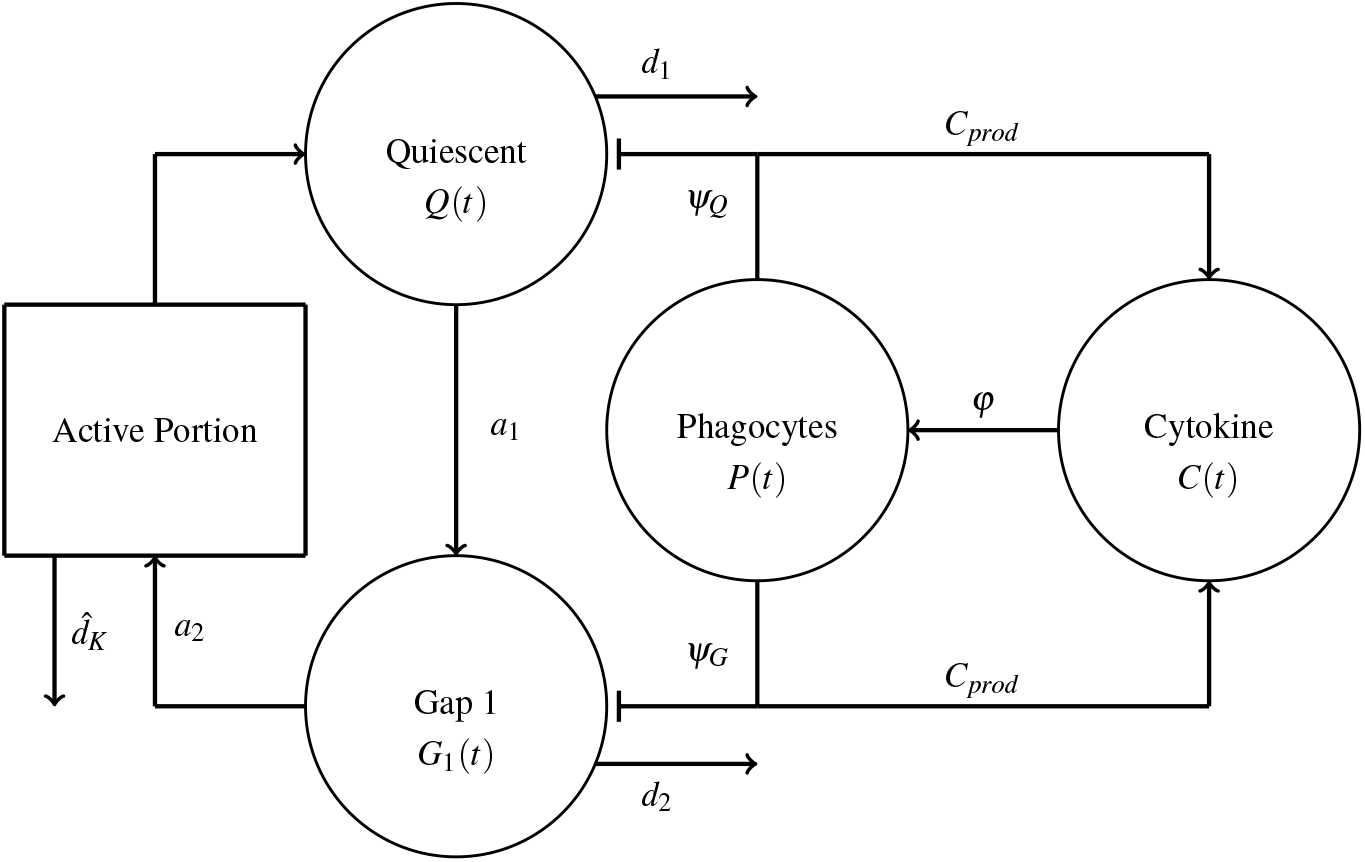
Pictorial representation of the tumour growth model. Populations are denoted by circles, processes by squares and rates by arrows. Quiescent cells enter *G*_1_(*t*) at rate *a*_1_ and undergo apoptosis at rate *d*_1_. Cells leave *G*_1_(*t*) and enter the active phase of the cell cycle at rate *a*_2_ while undergoing apoptosis at a rate *d*_2_. The active phase death rate is 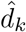 and cells re-enter quiescence after mitosis. Phagocytes interact with quiescent and *G*_1_ phase cells at respective rates *ψ_Q_* and *ψ_G_*. Tumour-immune interaction drives cytokine production through the function *C_prod_*.

### 2.1 Tumour Growth Model Development

RNA viruses replicate in infected cells during stages *G*_1_ through *M* of the Burns and Tannock [1970] model of the cell cycle. As previously noted, we separately model the quiescent (*Q*(*t*)) and *G*_1_ phase (*G*_1_(*t*)) tumour cell populations. Quiescent tumour cells undergo apoptosis at a rate *d*_1_. We denote the transit rate between the quiescent and *G*_1_ population as *a*_1_ Cells in *G*_1_ undergo apoptosis at a rate *d*_2_, and enter into the active phase of the cell cycle at a rate *a*_2_. We define the cell cycle duration as the time length of the active portion of the cell cycle, calculated as the time a cell takes between exiting *G*_1_ and re-entering *Q*.

We assume that the cell cycle time of tumour cells is a positive random variable with PDF *K*(*t*) satisfying

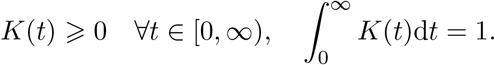

We assume that cells have an expected mean cell cycle duration of *τ*, so the expected value of *K*(*t*) satisfies

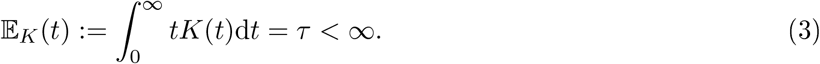

We will also use that

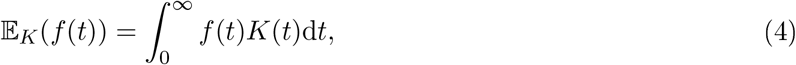
where in particular we note that the Laplace transform 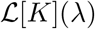 of the PDF *K*(*t*) is equivalent to 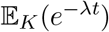 since

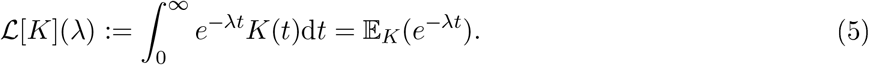

Let *A_R_*(*t*) denote the rate that successfully dividing cells re-enter quiescence at time *t*. Such cells began the active portion of the cell cycle some time σ in the past at rate *a*_2_*G*_1_(*σ*). The likelihood that these cells complete the cell cycle at time *t* is given by *K*(*t* − *σ*). Disregarding immune interaction for now, cells in the active portion of the cell cycle undergo apoptosis at a constant, distribution specific, rate 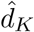. Consequently, cells that spend more time in the active phase of the cell cycle are more likely to undergo apoptosis instead of completing the cell cycle and returning to quiescence. Thus

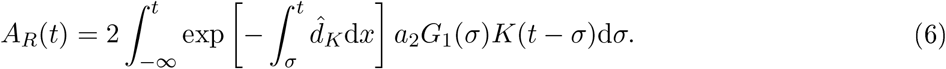

Later, we will update *A_R_*(*t*) to include tumour-immune interaction and viral therapy. The distributed delay expression *A_R_*(*t*) is a novel model of tumour cell reproduction which is more physiologically appropriate than a discrete delay.

The discrete delay model considered by Crivelli et al. [2012] corresponds to *K*(*t*) = *δ*(*t* − *τ*) and *d_δ_* = *d*_3_. The explicit link between equation (1) and the Crivelli model is shown in Appendix A. The expected cellular output of the cell cycle with a discrete and fixed duration is

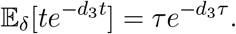

To ensure a consistent cellular output from the cell cycle for different distributions *K*(*t*), we define 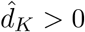 as the distribution dependent unique positive value that solves

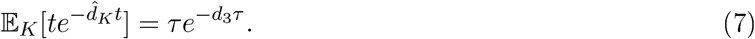

The parameter 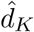 must exist for a given distribution *K* as the function

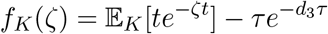
is continuous and satisfies

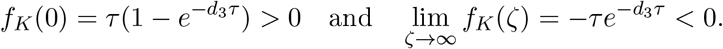

The intermediate value theorem along with the fact that *f*(*ζ*) is strictly decreasing for *ζ* > 0 guarantees the existence and uniqueness of 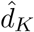

The resulting model of tumour growth without immunosurveillance is then

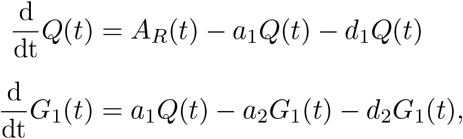
where *A_R_*(*t*) is given by (6).

### 2.2 Immune Model Development

The tumour microenvironment is complex and contains a multitude of cytokines and cell types [Bartlett et al., 2013; Cassady et al., 2016; Grivennikov and Karin, 2011; Hallam et al., 2009]. To avoid overcomplicating the model by adding variables and creating equations corresponding to each cytokine and signalling pathway, we instead model a general local proinflammatory cytokine compartment *C*(*t*). We assume the cytokine is produced at a variable rate *C_prod_*(*U*(*t*)) with the homeostatic production rate 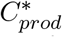. The viral and immune mediated destruction of tumour cells results in increased cytokine production by releasing tumour specific antigens [Bartlett et al., 2013; Bell and McFadden, 2014]. Conversely, we do not consider apoptosis of tumour cells to be immunogenic [Bartlett et al., 2013]. Therefore, *C_prod_*(*U*(*t*)) is an increasing function of viral and immune destruction of tumour cells. The resulting positive feedback loop is consistent with self activation of immune cells observed experimentally [Mosser, 2003]. Finally, we assume that the cytokine is cleared linearly at rate *k_elim_*, mimicking the dynamics of many endogeneous cytokines [Craig et al., 2016; Krzyzanski et al., 2010; Piscitelli et al., 1997]. The simplified cytokine dynamics are thus given by

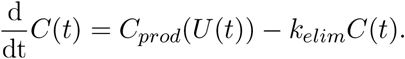

We assume that phagocytes can undergo phagocytosis multiple times, so phagocyte clearance is linear, and we do not include a phagocytosis related death term. Inflammatory cytokines drive phagocyte recruitment and activation [Bartlett et al., 2013; Cassady et al., 2016; Hallam et al., 2009]. Consequently, we model the local phagocyte population in a similar cytokine driven manner to Schirm et al. [2016] by using a Michaelis-Menten growth function *φ*(*C*(*t*)) with maximal production rate *k_cp_* and half effect concentration of cytokine *C*_1/2_. The phagocyte dynamics are therefore given by

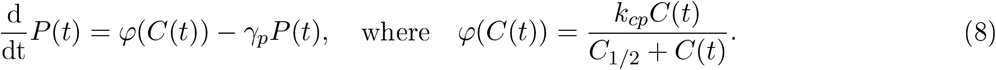

The disease free equilibrium concentrations of (*C*(*t*), *P*(*t*)) represent the tumour-free tissue concentrations of cytokine and phagocytes and are given by

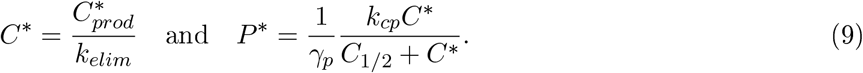

We describe phagocyte-tumour cell interaction by

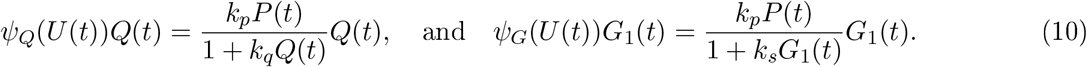

For small tumour cell populations, the tumour-immune interaction follows mass-action kinetics, while for large tumour cell populations, the phagocytosis rate is limited by the phagocyte concentration as would be expected. We assume that cells in the active portion of the cell cycle interact with the immune system in the same way as cells in the *G*_1_ phase.

The total immune mediated death is then

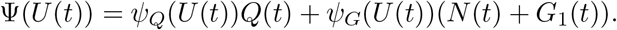

Contact rates similar to equation (10) were derived by Imran and Smith [2007] using a handling time argument.

### 2.3 Viral Therapy Model Development

Viral infections are caused by virus specific particles, called virions, that infect and replicate in host cells. Infected host cells die after undergoing lysis and releasing virions into the surrounding tissue. To model the effect of oncolytic virus treatment, we consider the virion population, *V*(*t*), and the number of infected malignant cells, *I*(*t*).

Infection occurs following contact of a virion and a susceptible cell. Susceptible cells are cells in the *G*_1_, *S, G*_2_ and *M* phases of the cell cycle. We model the infection rate between virions and susceptible cells by *η*(*U*(*t*)). Infection due to virion and susceptible cell contact occurs in a similar manner to tumour-immune interactions. Consequently, *η*(*U*(*t*)) is structured similarly to equation (10), with half effect concentration *η*_1/2_ and maximal infectious rate *κ*, so

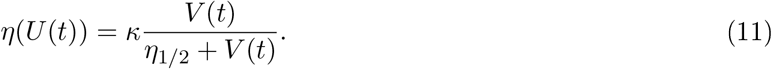

As previously noted, disease remission following viral therapy is thought to result from activation of the immune system against the tumour and increased antitumour immunity [Bartlett et al., 2013; Bell and McFadden, 2014; Cassady et al., 2016; Fukuhara et al., 2016; Rehman et al., 2016]. Therefore, introduction of viral therapy alone should not impact the stability of the disease free equilibrium but rather immune response to viral therapy may change the quantitative behaviour of solutions. This is in contrast to Crivelli et al. [2012], who modelled contact between virions and susceptible cells using a non-differentiable contact function. Their choice of contact function was motivated by noting that viral therapy has driven cancer into remission, which implicitly assumed that the virus alone drives disease remission.

Infected tumour cells are produced following infection and undergo lysis at a rate *δ*. Lysis of infected tumour cells releases *α* virions. Virions are only produced during lysis and lose infectivity at a rate *ω*, leading to the differential equations for *I*(*t*) and *V*(*t*)

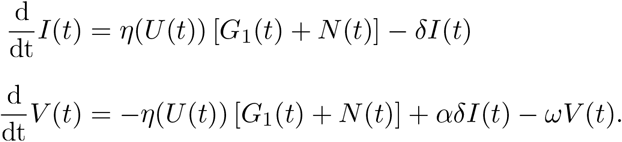

Clearance of proliferating cells leads to exponential loss as the cleared cells no longer divide nor return to quiescence. This is accounted for by updating equation (6) to include the loss of mitotic cells due to immune and viral mediated death, giving

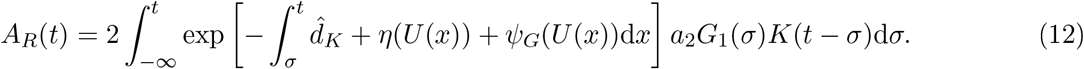

Finally, the link between the oncolytic virus and the immune system is cytokine production, modelled by *C_Prod_*(*U*(*t*)). Both lysis of infected cells and immune killing are immunogenic, leading to an increase in immune signalling. Therefore, we link virus and immune mediated cell death by the cytokine production rate *C_prod_*(*U*(*t*)), given by

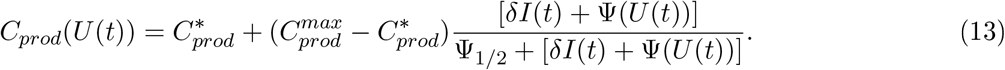

We note that 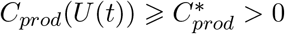 for nonnegative cell populations; the homeostatic cytokine production rate is effectively the minimal cytokine production rate.

Combining the differential equations for each population with the PDF *K*(*t*) gives the complete model in equation (1).

## 3 Model analysis

The mathematical model in equation (1) represents cell populations which are non-negative quantities. Consequently, we begin our analysis by showing that solutions of equation (1) evolving from non-negative initial data remain non-negative.

### Lemma 3.1.

*Assume that the parameters in equation* (1) *are strictly positive and that the initial conditions are componentwise non-negative. Moreover, assume that G*_1_(*s*) = *ϕ_G_*(*S*) ≥ 0 *for s* ∈ (−∞, 0]. *Then solutions of the initial value problem corresponding to equation* (1) *are non-negative for all time t* ≥ 0.

*Proof*. By the assumption on the initial conditions, 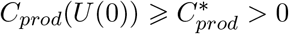, so

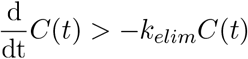
in a neighbourhood *t* ∈ [0, *ε_C_*]. Gronwall’s inequality ensures that 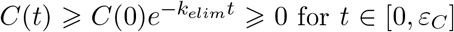. In this interval,

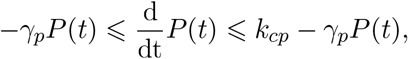
therefore

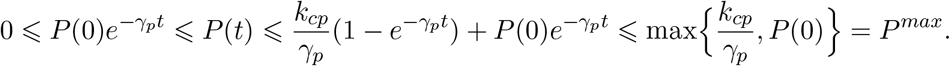

We now investigate the populations *Q*(*t*) and *G*_1_(*t*). If *Q*(0) = *G*_1_(0) = 0 and *ϕ_G_*(*s*) = 0 *K*-almost everywhere in (−∞, 0], *Q*(*t*) and *G*_1_(*t*) remain identically zero for all time *t* > 0 If *Q*(0) = 0 and *ϕ*(*s*)*K*(−*s*) > 0 on some set of positive measure in (−∞ 0], then *Q*(*t*) eventually becomes positive for some *t* > 0. Therefore, we only need to consider the case where *Q*(0) > 0 and *ϕ*(*s*) ≥ 0 for *s* ∈ (−∞, 0].

Now, let *t_g_* ∈ [0, *ε_C_*] be the first time that *G*_1_(*t_g_*) = 0. Then *A_R_*(*t*) defined by equation (12) satisfies *A_R_*(*t*) ≥ 0 for all *t* ∈ [0, *t_g_*]. It follows from equation (1) that

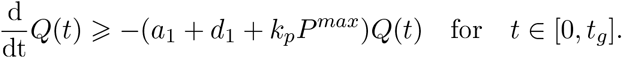

Then *Q*(*t*) > 0 for *t* ∈ [0, *t_g_*] and

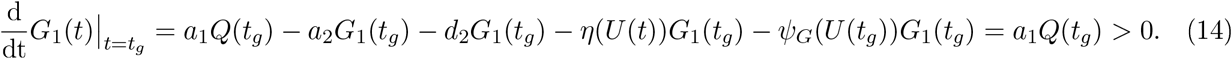

Thus *G*_1_(*t*) is strictly increasing at *t_g_*. If *t_g_* = 0, then *G*_1_(*t*) > 0 immediately. Conversely, if *t_g_* > 0, then *G*_1_(*t*) must be nonincreasing at *t_g_*. This contradicts equation (14), so no such *t_g_* > 0 can exist and *G*_1_(*t*) > 0 for *t* ∈ (0, *ε_C_*]. Since *A_R_*(*t*) ≥ 0 while *G*_1_(*t*) ≥ 0 it follows from the arguments above that *Q*(*t*) > 0 while *G*_1_(*t*) ≥ 0. Finally, it is simple to see that *G*_1_(*t*) > 0 for *t* ∈ (0, *ε_C_*] implies that *N*(*t*) defined by (2) satisfies *N*(*t*) > 0 for all *t* ∈ (0, *ε_C_*].

If *V*(0) = *I*(0) = 0 then the *I*(*t*), *V*(*t*) populations remain identically zero for all time. Therefore, we consider *V*(0) + *I*(0) > 0 and we have three cases:

**Case I** If *V*(0) = 0 then *I*(0) > 0 and it is simple to calculate that

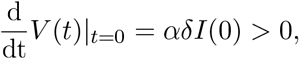
so V (*t*) becomes strictly positive immediately.

**Case II** If *I*(0) = 0

If *Q*(0) = 0 and *ϕ*(*s*) = 0 almost everywhere in (−∞, 0], the tumour free case, then *Q*(*t*), *G*_1_(*t*) and *I*(*t*) remain identically zero for all time *t* > 0 and *V*(*t*) decays exponentially to 0.

Thus, as above, we need only consider *Q*(0) > 0 and *G*_1_(*t*) > 0 in (0, *ε_C_*]. Now, *I*(0) = 0 so *V*(0) > 0 and for all *t* ∈ (0, *ε_C_*], if *I*(*t*) = 0 then

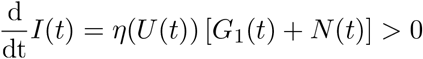
and *I*(*t*) > 0 for all *t* ∈ (0, *ε_C_*], otherwise a contradiction ensues.

**Case III** Thus, it only remains to consider the case where *V*(*t*) and *I*(*t*) are both strictly positive immediately and remain positive in some neighbourhood of *t* = 0. While *I*(*t*) and *V*(*t*) are non-negative, we compute

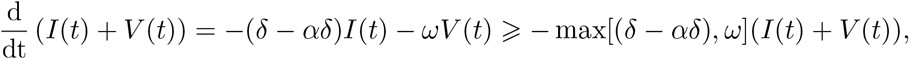
so 
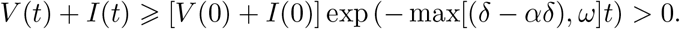

If there exists a time *t_v_* such that *V*(*t_v_*) = 0 then 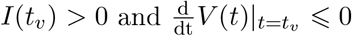, but arguing as in Case I, we see that 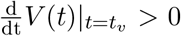, and hence no such time *t_v_* can exist. Similarly, if there exists a time *t_I_* such that *I*(*t_I_*) = 0 then 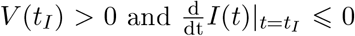, but arguing as in Case II, we see that 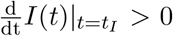, so no such *t_I_* can exist. Therefore, *V*(*t*) > 0 and *I*(*t*) > 0 for all *t* ∈ [0, *ε_C_*].

Finally, for *Q*(*t*), *G*_1_(*t*) *I*(*t*), *V*(*t*), *P*(*t*) strictly positive, the cytokine production rate satisfies *C_prod_*(*U*(*t*)) ≥ 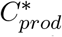, so

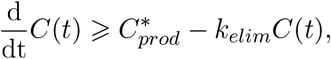
and 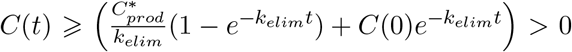 for all *t* ∈ [0, *ε_C_*]. Then, each component is positive at *t* = *ε_C_* and the above argument extends from [0, *ε_C_*] to [0, ∞).

### 3.1 Linearisation of the distributed DDE

The system (1) has the cancer free equilibrium (CFE), *U*\* = (0, 0, 0, 0, *C*\*, *P*\*). Although it is often convenient to regard a trajectory *U*(*t*) of the system (1) as a parameterised curve with with 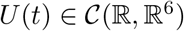, it is important to realise that the DDE system (1) defines an infinite dimensional dynamical system. The infinite-dimensional phase space is

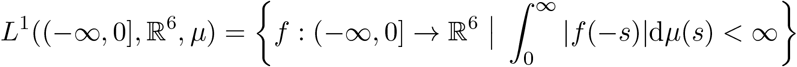
where | · | is the *ℓ*_1_ norm in 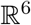, and *μ* is a probability measure whose Radon-Nikodym derivative with respect to the Lebesgue measure is *K*(*t*). When *K*(*t*) is Riemann integrable (such as in the case of the Gamma distribution that we will consider in Section 4) this implies that

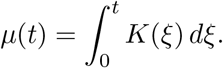

This space satisfies the axioms given by Hale and Verduyn Lunel [1993]; Hino et al. [1991], so there exists a unique solution to the corresponding initial value problem.

To investigate the long term behaviour of the model, we linearise the system around the CFE in *L*_1_(*μ*). In a similar procedure to Câmara De Souza et al. [2018], we first linearise the function *A_R_*(*t*), given in equation (12) around the CFE. Using the Taylor expansions of *η*(*U*(*x*)) and *ψ_G_*(*U*(*x*)), with *η*(*U*\*) = 0, we approximate the inner integral

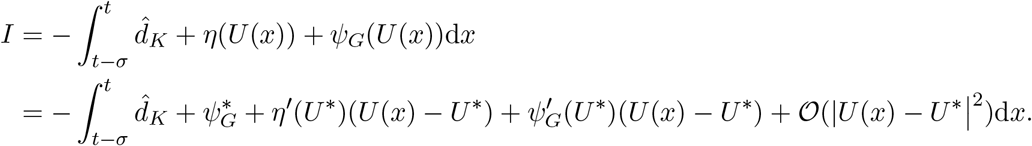

The full expansion of *e^I^* is

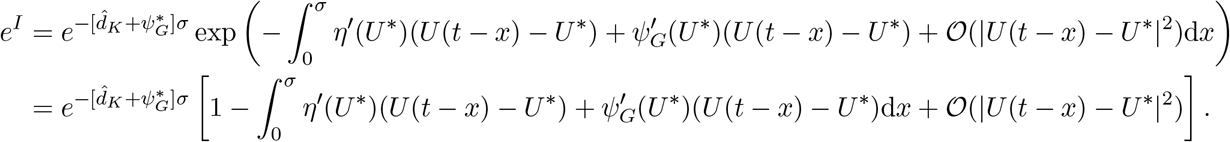

Importantly, *e^I^* is multiplied by *G*_1_(*t* − σ) in *A_R_*(*t*) and any non-constant terms of *U*(*t*) in the expansion of *e^I^* are consequently nonlinear. So we obtain

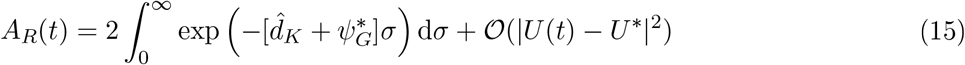

We translate the CFE of equation (1) to zero by setting 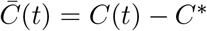 and 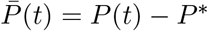 with *C*\* and *P*\* given by equation (9). Then, noting that *η*(*U*\*) = 0 and using (15), the *N*(*t*) terms in the *I*(*t*) and *V*(*t*) equations are also nonlinear. Equation (1) becomes

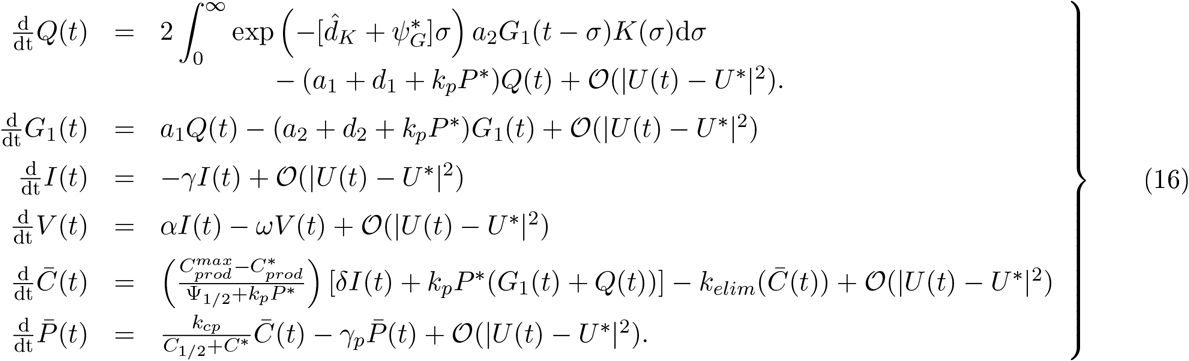

We follow Smith [2011] to complete the linearisation. We define **X**(*t*): = *U*(*t*) — *U*\* and use **X**_*τ*_ to denote the linear delayed terms via

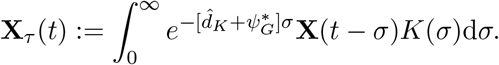

By making the ansatz **X**(*t*) = **C***e^λt^*, we see that **X**_*τ*_(*t*) satisfies

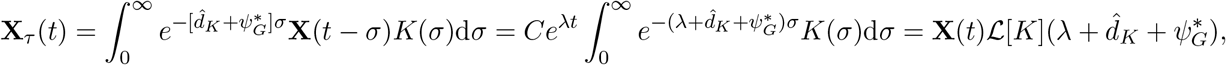
where 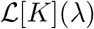 is the Laplace transform of *K*(*σ*) defined by (5).

Dropping the nonlinear terms in equation (16) and setting

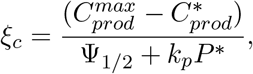
we obtain the linearised infinite dimensional DDE

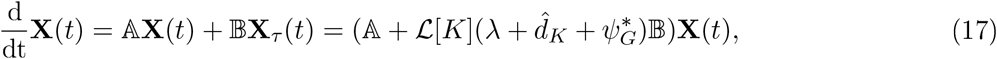
where

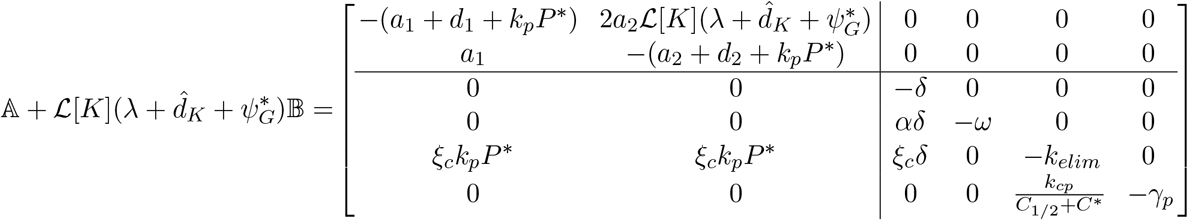

Hence equation (17) becomes

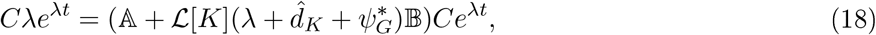

From equation (18), the characteristic equation is

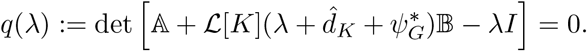

Using the block nature of the linearisation matrix gives

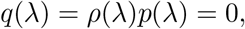
where

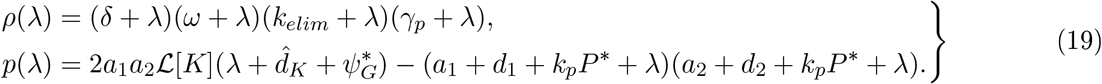

Here *ρ*(*λ*) is the determinant of the lower triangular block and has strictly negative real roots. The explicit roots of *ρ*(*λ*) imply that the stability of the CFE is determined by the roots of *ρ*(*λ*).

To study the persistence of small tumours, we characterise the stability of the disease free steady state. Typically, for DDEs, this involves solving a transcendental equation with infinitely many roots. To simplify the following analysis, we first show that the rightmost root of the characteristic equation is real. This result is unsurprising, as a complex rightmost eigenvalue would give rise to spiralling solutions around the CFE, which would become negative, contradicting **Lemma 3.1.**

#### Lemma 3.2.

*For strictly positive parameters, the rightmost root of q*(*λ*) *is real*.

*Proof*. First, we note from (5) that the Laplace transform of a non-negative function *f*, is a decreasing function of *λ*. Similarly,

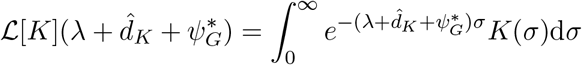
is decreasing for real *λ* where it converges. Therefore, as a function of a real variable, *p*(*λ*) is continuous and *p*(*λ*) is strictly decreasing for

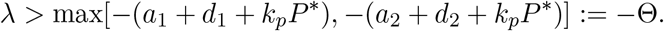

Moreover,

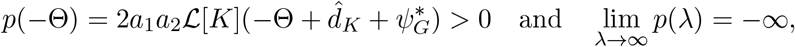
so there is exactly one real root *λ*\* of *p*(*λ*) that satisfies *λ*\* > −Θ.

Since *ρ*(*λ*) has strictly negative real roots, any complex roots, *v* = *v_r_* + *iv_i_* with *v_r_* ∈ (−Θ, ∞) and *v_i_* ≠ 0, of the characteristic equation *q*(*λ*) must solve *p*(*v*) = 0, which we may rewrite as

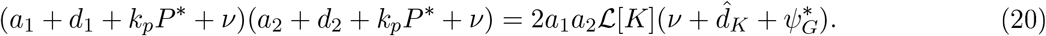

Taking the magnitude of the equality (20) gives

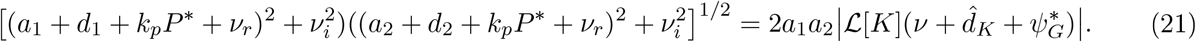

However,

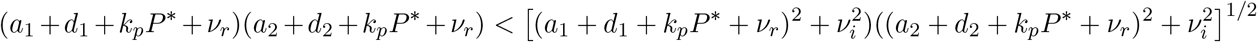
and

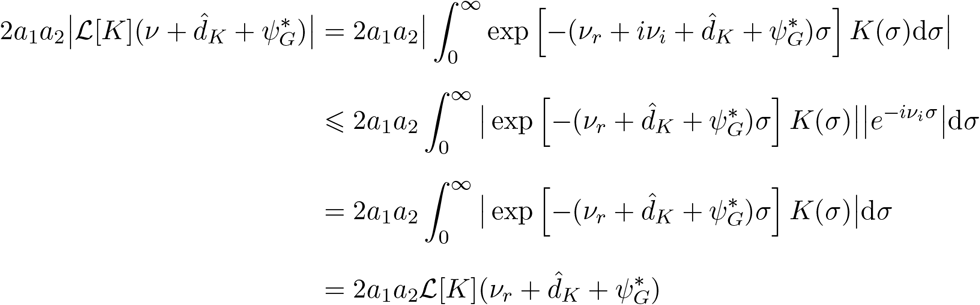
where the last equality comes from the nonegativity of the integrand. Substituting these bounds into equation (21) gives

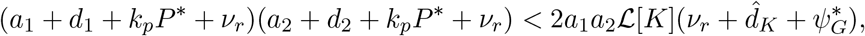
from which we obtain

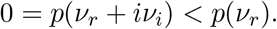

Since *p*(*λ*) is strictly decreasing for *λ* > −Θ, we must have *v_r_* < *λ*\*. Then, the rightmost root of *q*(*λ*) is either *λ*\* or a root of *ρ*(*λ*) and is real.

The preceding result simplifies the analysis of the transcendental characteristic equation by ensuring that the critical characteristic root is real. Therefore, the stability of the CFE, and consequently, the persistence of small tumours, can be characterised using the intermediate value theorem.

#### Theorem 3.3.

*The cancer free equilibrium, U* of equation* (1) *is locally stable if*

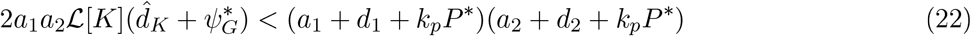
*and unstable if*

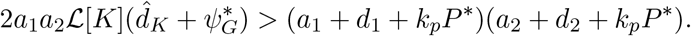

*Proof*. The condition for stability is equivalent to *p*(0) < 0. In this case, since *p*(*λ*) is strictly decreasing for *λ* > max[−(*a*_1_ + *d*_1_ + *k_p_P*\*), − (*a*_2_ + *d*_2_ + *k_p_P*\*)], there can be no real root of the characteristic equation with non-negative real part. Since the rightmost root must be real, all roots of the characteristic equation must have negative real part and the CFE is stable.

The condition for instability is equivalent to *p*(0) > 0. Since

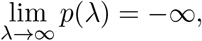
the intermediate value theorem ensures that there is a root of the characteristic equation in the positive half plane and the CFE is unstable. □

Using (5) we can rewrite the stability condition (22) as

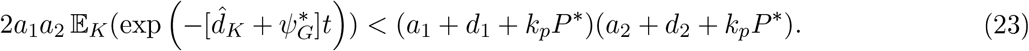

This can be rearranged as a basic reproduction number type condition

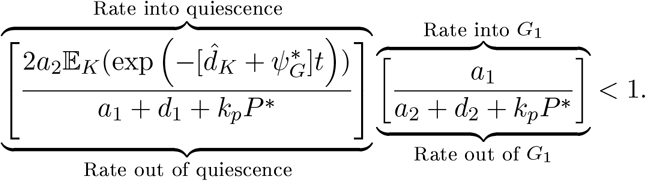

Hence, the CFE is locally attracting if the product of the ratios of expected transit rates into and out of the quiescent and *G*_1_ phases is less than one. Biologically, this corresponds to each cell that transits out of either the quiescence or *G*_1_ phase not replacing itself through mitosis.

Finally, we can characterise the importance of heterogeneity in cell cycle duration as a determining factor of disease progression. Let 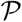 be the parameter space of the distributed DDE (1). Following Campbell and Jessop [2009], for each PDF *K*(*t*), we define the stability region as

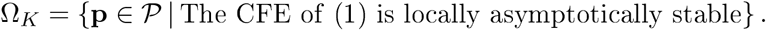

Then, we are able to characterise the stability regions for certain PDFs with respect to the discrete DDE. For these PDFs, the tumour heterogeneity in cell cycle duration acts to destabilise the CFE and leads to more a robust tumour. We formalise this relationship in the following corollary.

**Corollary 3.4.** *For any PDF K*(*t*) *which satisfies* (3) *and* 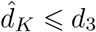 *we have the indusion* Ω_*K*_ ⊆ Ω_*δ*_.

*Proof*. Take **p** ∈ Ω_*K*_ so that equation (23) is satisfied and the CFE is locally stable. Now, we define

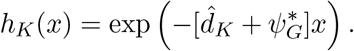

It is simple to see that *h_K_*(*x*) is convex. Jensen’s inequality gives

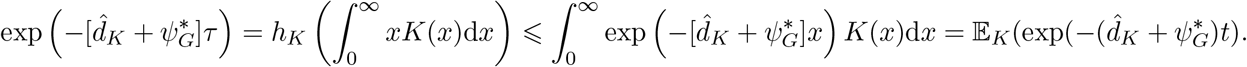

Now, using 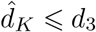, we have

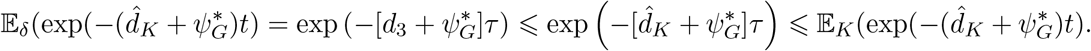

It follows that

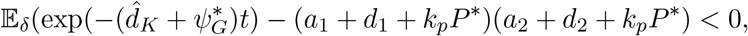
so the CFE is stable in the discrete DDE case and **p** ∈ Ω_*δ*_.

The condition 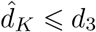 corresponds to

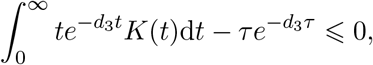
which can be viewed as a measure of the skewness of the PDF *K*(*t*). Using equation (3), this condition is satisfied if

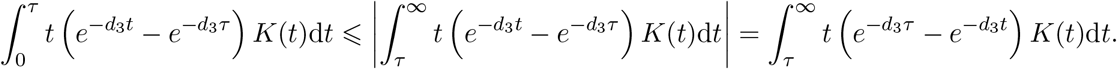

It is important to note that the linearisation only determines local stability. So, while small tumours may not grow, large tumours do not necessarily disappear. In fact, for a given level of immune recognition of tumour cells, *k_p_*, there is a critical tumour size above which the tumour grows unboundedly. The critical tumour size acts as a separatrix between tumour extinction and growth and takes the form of a nonzero equilibrium point where tumour growth and immune surveillance are balanced. In Theorem 3.5, we show that such an equilibrium must exist. Transition across this equilibrium has been hypothesised to occur as part of the cancer immunoediting process that allows tumours to grow and corresponds to a transient decrease of *k_p_* [Bhatia and Kumar, 2011; Mittal et al., 2014; Swann and Smyth, 2007].

To emphasise the biological interpretation of Theorem 3.5, we use the stability condition as written in equation (23) to characterise the existence of the non-zero equilibrium.

#### Theorem 3.5.

*Assume that the parameters in equation* (1) *are nonnegative. Let* 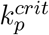 *solve*

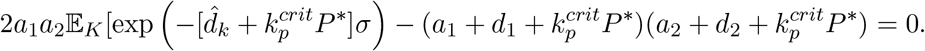

*Then, for* 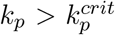*, there exists a strictly positive untreated equilibrium solution* 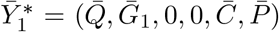 *of equation* (1) *with Q*_1_ *and G*_1_ *strictly positive*.

*Proof*. First, in the absence of viral treatment, *V*(0) = 0 and *I*(0) = 0, so (*V*\*, *I*\*) = (0, 0).

To simplify notation in the proof, we set *ξ*_*i*_ = *a_i_* + *d_i_* + *k_p_P*\* for *i* = 1, 2. We consider the differential equation for *G*_1_(*t*) at equilibrium, so 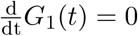 and

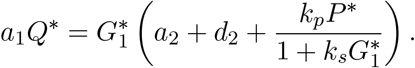

This can be rearranged as a quadratic equation in 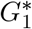

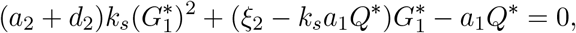
whose positive root is a function of *Q*\* defined by

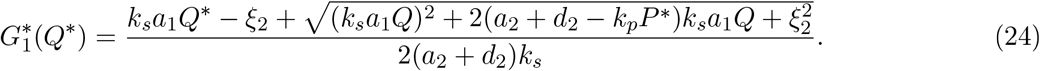

Now, inserting 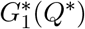 into 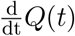 gives

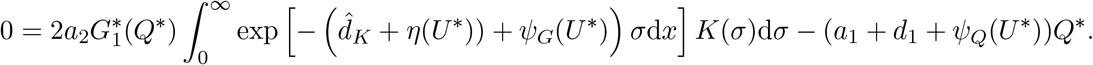

Using (4) gives

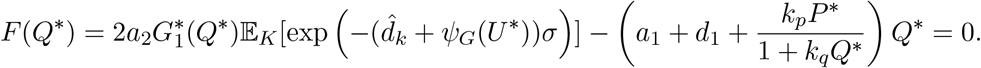

We write

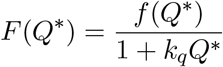
where

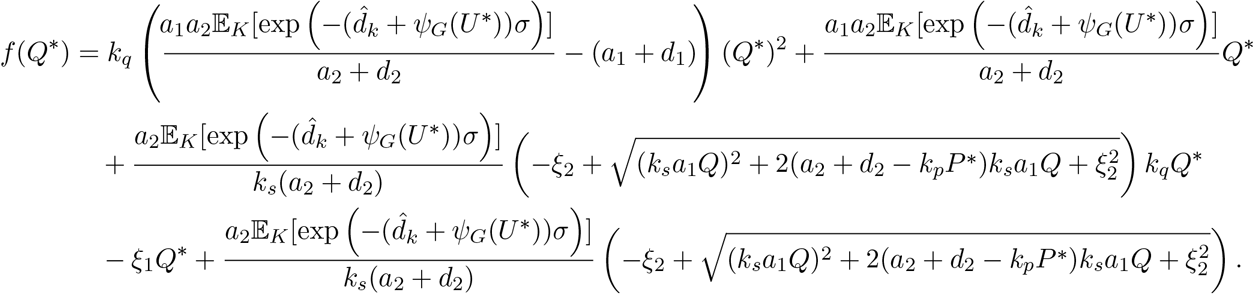

The equilibrium concentration *Q*_1_ must therefore solve *f*(*Q*_1_) = 0. A simple calculation shows that *f*(0) = 0, so we search for *Q*_1_ positive. Now, as *Q*\* → ∞.

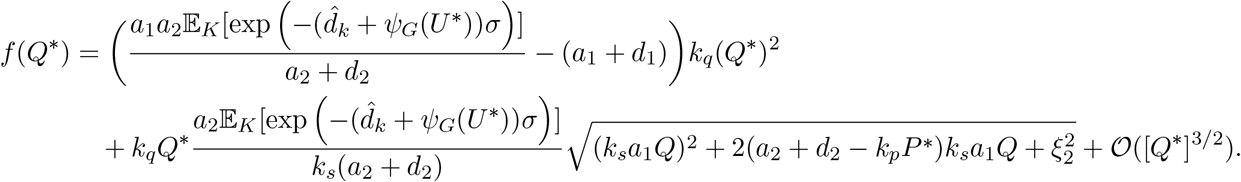

This is equivalent to

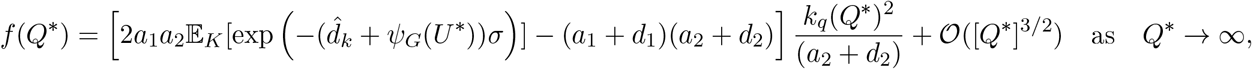
so the sign of 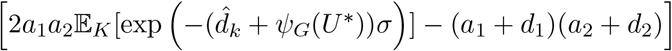 determines the sign of *f*(*Q*\*) as *Q*\* grows infinitely large. Now,

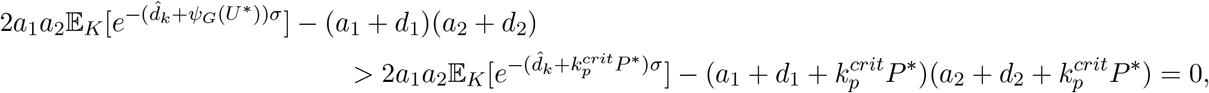
so *f*(*Q*\*) grows infinitely large with *Q*\* and must be positive for large values of *Q*\*.

Next, as *Q*\* → 0,

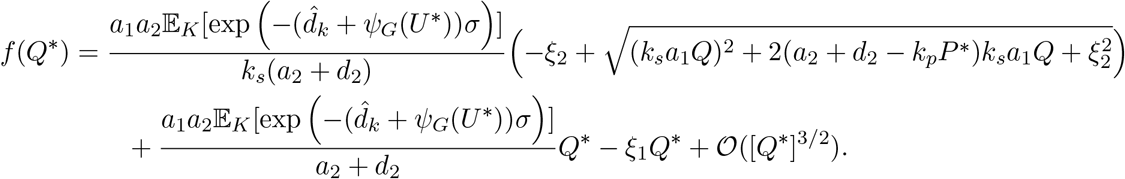

Taylor expanding the square root about the point *Q*\* = 0 gives

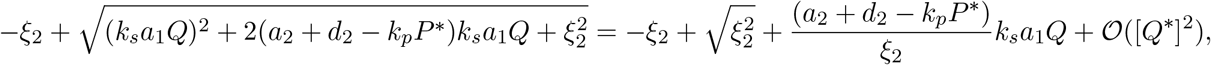
so for *Q*\* near 0,

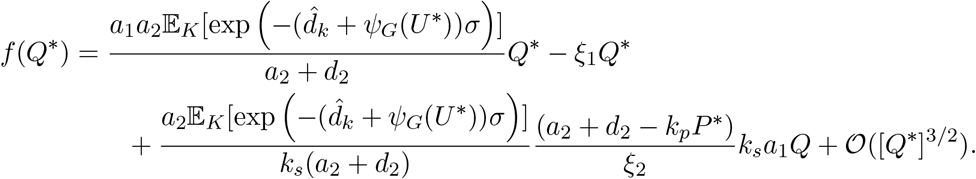

Crucially, *a*_2_ + *d*_2_ − *k_p_P*\* = 2(*a*_2_ + *d*_2_) − *ξ*_2_, so with *f*(0) = 0

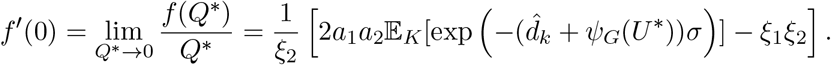

Thus, the sign of *f*′(0) is determined by the sign of

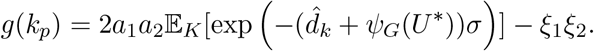

The function *g*(*k_p_*) is strictly decreasing with 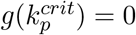, therefore, *f*′(0) < 0 for 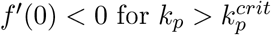.

Consequently, *f*(*Q*\*) is negative for *Q*\* small and positive, and positive for large *Q*\*, so there must be a positive root 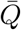 with 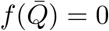. This root defines a solution 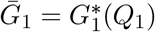 of equation (24).

Finally, we can write an equilibrium solution of 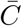 as a function of *C*(*t*) via

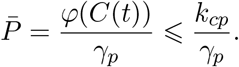

Given the upper bound of 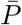 and the pair 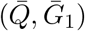, the function Ψ(*U*(*t*)) is bounded. Therefore, there must exist a solution 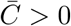 to

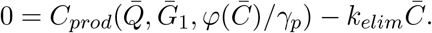

Finally, using the value of 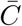, we can calculate the corresponding equilibrium 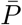. □

## 4 The Gamma Distribution and Equivalent ODE System

To translate our results for a generic distribution into predictions of tumour growth, we must specify a distribution of cell cycle durations, corresponding PDF *K*(*t*), and death rate 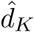. We assume that cell cycle durations follow a gamma distribution, so 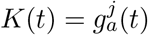. The function 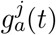 is the PDF of the gamma distribution with

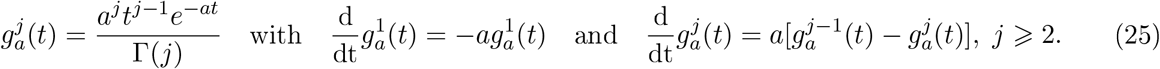

The real positive parameters *a* and *j* in equation (25) define the shape of the gamma distribution. The expected cell cycle duration is *τ* = *j*/*a*. For given *τ* we take *j* to be a strictly positive integer and determine *a* by *a* = *j*/*τ*. The standard deviation, *s*^2^, of the gamma distribution is given by *s*^2^ = *τ*^2^/*j* For fixed *τ*, larger values of *j* result in a more concentrated distribution about *τ*. In Appendix A we demonstrate that in the limit as *j* → ∞ (with fixed *τ*) the gamma distributed model converges in distribution to a delta distributed model with discrete delay *τ*.

To calculate 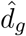, we note that the expected cellular output of the cell cycle is

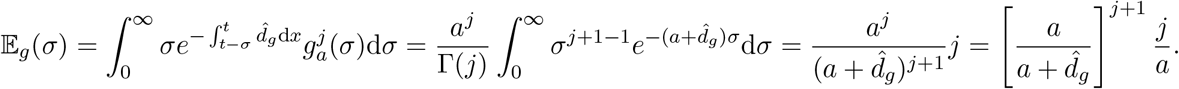

Imposing the equality (7) and *τ* = *j*/*a* gives

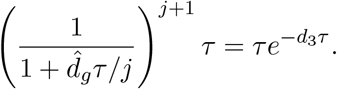

Therefore, 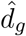 is given by

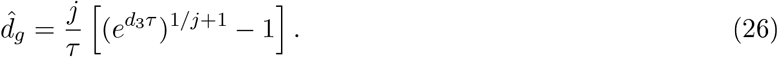

### 4.1 Equivalent ODE Formulation

The link between gamma distributed DDEs and transit chain ODEs has been known since at least the 1960s [Vogel, 1961]. The equivalence between infinite dimensional DDEs and ODEs is typically established through the linear chain technique. Among many other areas, the linear chain technique has recently been used in the pharmaceutical sciences [Câmara De Souza et al., 2018; Hu et al., 2018]. More generally, the equivalence between distributed DDEs and ODEs was studied by Diekmann et al. [2017].

Typical applications of the linear chain technique involve a transit chain type ODE without growth or loss throughout the chain. In the example of cellular growth represented by a transit chain model, the number of cells is conserved throughout the delayed process. Here, we derive a variant of the linear chain technique that accounts for the exponential decay of the mitotic cell population due to apoptosis, immune pressure and lysis as modelled in equation (1). The resulting ODE system is a compartment model with linear clearance throughout the transit chain.

By taking 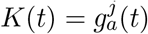 with 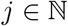 and *a* = *k_tr_* = *j*/*τ* and setting

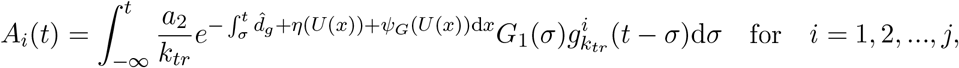
we can reduce the distributed DDE model to a system of ODEs. We show in Theorem 4.2 that equation (1) is equivalent to the system of ODEs

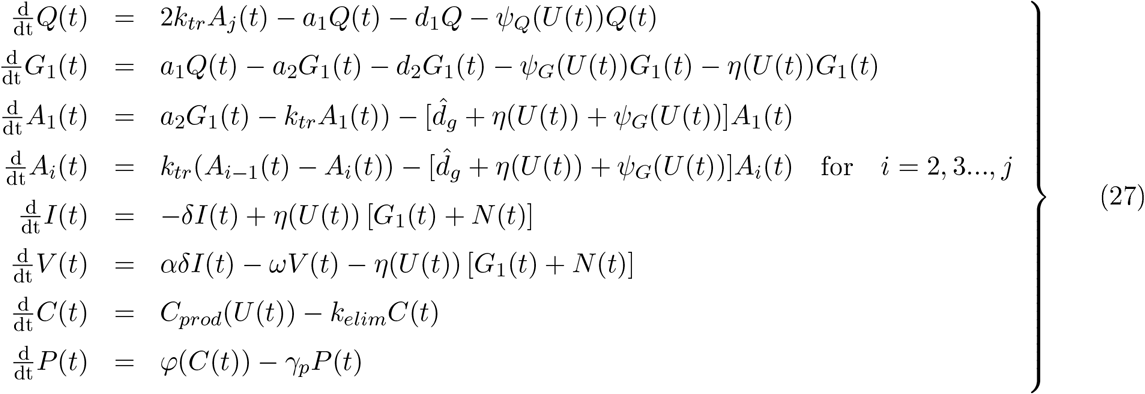
with identical initial conditions to the distributed DDE for *Q*(0), *V*(0), *I*(0), *P*(0), *C*(0) and

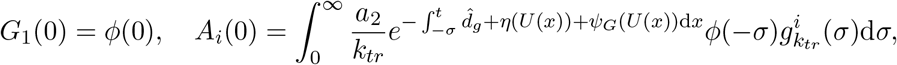
where ϕ(*s*) is the history function of equation (1).

#### Lemma 4.1.

For an integrable function *G*_1_(*t*) *and* 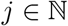 *with a = k_tr_ = j/τ, the vector with i-th component given by*

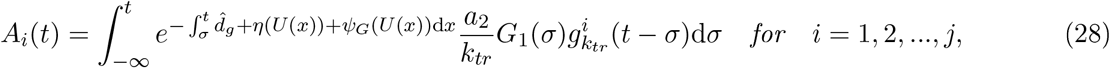
*is the solution of the system, of differential equations given by*

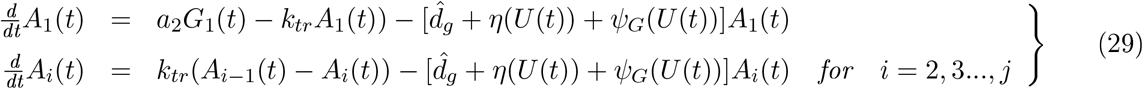

*Proof*. Using the Lebniz and product rules, we differentiate *A*_1_(*t*) to obtain

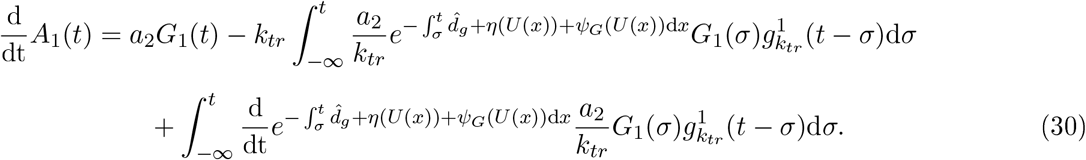

Computing the derivative of the exponential then gives

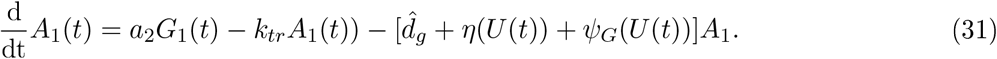

Similarly for general *i*, differentiating the expression for *A_i_*(*t*) from (28) gives

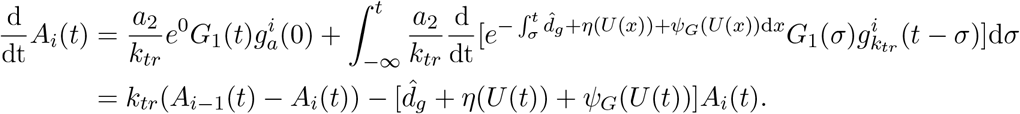

Thus, the vector **A**(*t*) = [*A*_1_(*t*), *A*_2_(*t*), …, *A_j_*(*t*)] satisfies equation (29). □

Comparing equations (30) and (31) shows that the exponential loss of cells during the cell cycle in equation (1) corresponds to linear clearance in the equivalent transit compartment system of ODEs.

We now show the equivalence of the ODE and DDE models by using **Lemma 4.1** to replace the integral terms in equation (1).

#### Theorem 4.2.

*The system of distributed DDEs* (1) *with* 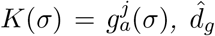 *as given in* (26) *and initial conditions Q*(0) = *Q*0, *I*(0) = *I*0, *C*(0) = *C*_0_ *and history functions V*(*s*) = *ϕ_V_*(*s*), *P*(*s*) = *ϕ_P_*(*s*) *and G*_1_(*s*) = *ϕ_G_*(*s*) *for s* ∈ (−∞), 0] *is equivalent to the system of ODEs* (27) *with initial conditions Q*(0) = *Q*_0_, *I*(0) = *I*_0_, *C*(0) = *C*_0_, *V*(0) = *ϕ_V_*(0), *P*(0) = *ϕ_P_*(0), *G*_1_(0) = *ϕ_G_*(0) *and*

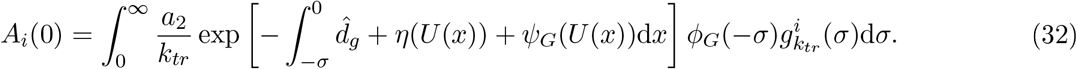

*Proof*. Using **Lemma 4.1**, we see that

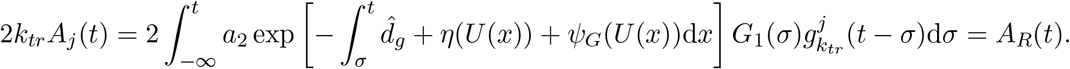

Thus, the differential equations for *Q*(*t*) in (1) and (27) are equivalent.

The remaining terms in equation (27) are exactly those in equation (1). To finish the conversion from the DDE (1) to the ODE (27), we must specify the initial conditions. Given the history functions [*ϕ_G_*(*s*),*ϕ_V_*(*s*), *ϕ_P_*(*s*)] from the DDE model, we chose the initial conditions *A_i_*(0) of equation (27) according to equation (32). This ensures that the solution of equation (27) is equivalent to the solution of equation (1) [Smith, 2011].

To convert from the ODE (27) to the DDE (1), we must take care with the construction of the history functions (*ϕ_G_*(*s*),*ϕ_V_*(*s*), *ϕ_P_*(*s*). The ODE is equipped with initial conditions *V*(0) and *P*(0). For simplicity, we set *ϕ_V_*(*s*) = *V*(0) and *ϕ_P_*(*s*) = *P*(0).

The *j* initial conditions for each *A_i_*(0) define *j* constraints on *ϕ_G_*(*s*). There are many history function that satisfy these constraints and the ODE reduction of the DDE defines the same solution for each such history function. We show how to construct one such history function *ϕ_G_* ∈ *L*^1^((−∞, 0], 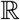, *μ*). Let the ODE system have initial conditions

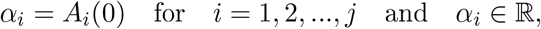
and chose a sequence of points

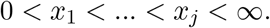

Now, we make the following ansatz for *ϕ_G_*(*S*)

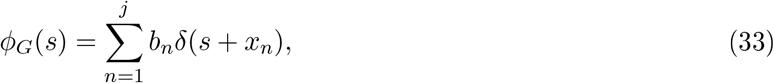
where *δ*(*x*) is the Dirac function. We will show that is possible to chose the 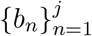 such that

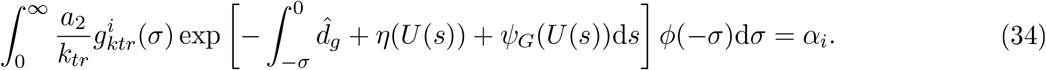

However, the histories *ϕ_v_* (*s*), *ϕ_P_*(*s*) and *ϕ_G_*(*s*) appear in the integral term

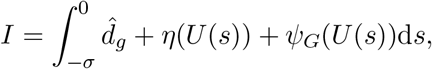
so some care is needed. We have already set *ϕ_V_*(*s*) = *V*(0) so *η*(*U*(*s*) is defined on (−∞, 0], so we need only consider

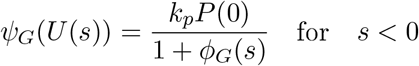
with *ϕ_P_*(*s*) = *P*(0) Inserting equation (33) for *ϕ_G_*(*s*) gives

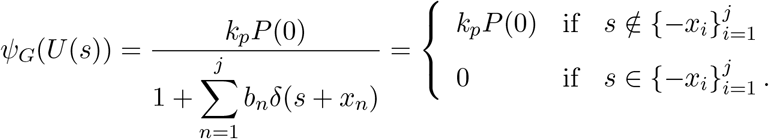

Since *ψ_G_* only appears in a Lebesgue integral and differs from *k_P_P*(0) on a set of measure 0, the following holds 
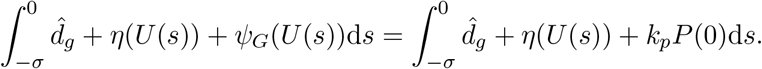

Therefore, finding 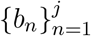 such that equation (34) holds is equivalent to finding 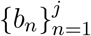 such that

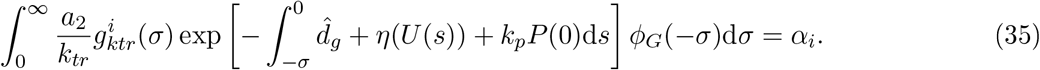

Using the ansatz for *ϕ_G_* in equation (35) gives the following system of equations for *i* = 1, 2, …, *j*

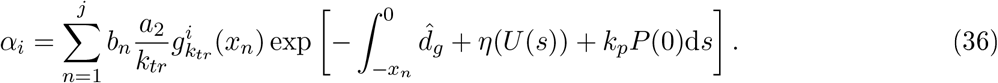

To simplify notation, set

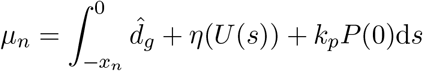
and note *μ_n_* is independent of the unknowns 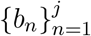.

Equation (36) defines a linear system of equations for the unknowns 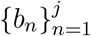. Consequently, there exists a unique solution to (36) if the matrix

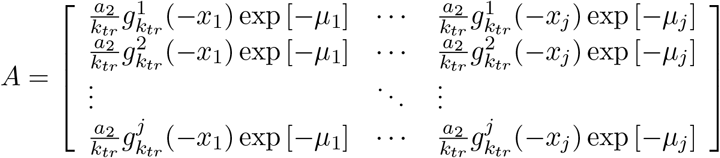
is invertible. To show this matrix is invertible, we will show that det(*A*) ≠ 0. Using the definition of 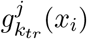. the *m*-th column has a common factor of

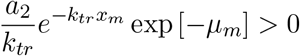
while, the *n*-th row has a common factor of 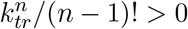 for *n, m* = 1, 2,.., *j*. Thus

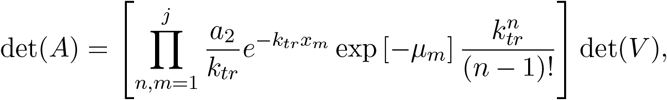
where 
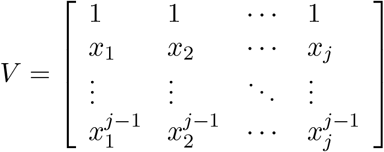

Since *V* is a Vandermonde Matrix and the 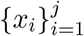 are distinct, det(*V*) ≠ 0. Consequently, det(*A*) ≠ 0 so *A* is invertible and we can uniquely determine the 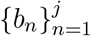. □

The equivalence between ODEs and gamma distributed DDEs has been used extensively since Vogel [1961]. Some authors have shown how to convert ODE transit compartment models to distributed DDE for specific initial conditions [Câmara De Souza et al., 2018; Cooke and Grossman, 1982]. However, to the author’s knowledge this is the first proof of direct equivalence between an ODE and a distributed DDE for arbitrary ODE initial conditions established by explicitly constructing a suitable history function.

### 4.2 Numerical Results

For the purpose of numerical simulation, the system of finite dimensional ODEs derived in Section 4.1 is much more tractable than the distributed DDE. Numerically solving the distributed DDE requires the development and implementation of a numerical differential equation solver capable of accurately computing the semi-infinite convolution integral, while there are numerous existing methods for solving systems of ODEs. To solve the DDE given in equation (1), we simulate the equivalent ODE in equation (27) and calculate *N*(*t*) as shown in Appendix B to illustrate the analytical results of Section 3.

For simplicity, we only present the dynamics of *Q*(*t*), as these dynamics are representative of the full model’s behaviour. The parameters used in these simulations are given in Table 1.

**Table 1:** The parameters used to simulate equation (27) inFigure 3. *C*_1/2_ was calculated from the homeostatic phagocyte production rate and *k_p_* was calculated from the mass-action tumour-immune interaction from Liu et al. [2007]. 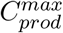 was calculated from G-CSF response to infection [Pauksen et al., 1994]. *η*_1/2_ was chosen to ensure a high initial infectivitv of viral therapy while *k_q,s_* and Ψ_1/2_ were selected to give physiologically realistic simulations.

| Parameter | Value | Biological Interpretation (Unit) | Reference |
| --- | --- | --- | --- |
| $a_1$ | 0.9 | Quiescent to interphase rate (1/day) | Crivelli et al. [2012] |
| $d_1$ | $1 \times 10^{-5}$ | Quiescent death rate (1/day) | Crivelli et al. [2012] |
| $a_2$ | 0.7 | Interphase to active phase rate (1/day) | Crivelli et al. [2012] |
| $d_2$ | 0.19 | Interphase death rate (1/day) | Crivelli et al. [2012] |
| $d_3$ | 0.19 | Active phase death rate (1/day) | Crivelli et al. [2012] |
| $\hat{d}_g$ | 0.167 | Active phase death rate (1/day) | Calculated from equation (26) |
| $\kappa$ | 1.15 | Virion contact rate (1/day) | Crivelli et al. [2012] |
| $\eta_{1/2}$ | $V(0)/10$ | Virion half effect concentration (virions) | See caption |
| $\delta$ | 1.119 | Lysis rate (1/day) | Crivelli et al. [2012] |
| $\alpha$ | 1.65 | Lytic virion release rate (virions/cell) | Crivelli et al. [2012] |
| $\omega$ | 0.75 | Virion death rate (1/day) | Crivelli et al. [2012] |
| $k_{cp}$ | 6.63 | Maximal phagocyte production rate ($10^{10}$ cells/day) | Schirm et al. [2016] |
| $C_{1/2}$ | 0.87743 | Phagocyte production half effect (ng/mL/day) | Liu et al. [2007] |
| $\Psi_{1/2}$ | 7 | Cytokine production half effect ($10^{10}$ cells/day) | See caption |
| $\gamma_p$ | 1 | Phagocyte death rate (1/day) | Liu et al. [2007] |
| $C_{prod}^*$ | 0.014161 | Homeostatic cytokine production rate (ng/mL/day) | Craig et al. [2016] |
| $C_{prod}^{max}$ | 1.4161 | Maximal cytokine production rate (ng/mL/day) | See caption |
| $k_{elim}$ | 0.16139 | Cytokine elimination rate (1/day) | Craig et al. [2016] |
| $k_p$ | 0.065 | Phagocyte-tumour cell contact rate (1/day) | Liu et al. [2007] |
| $k_{q,s}$ | 1.75 | Phagocyte cell digestion constant | See caption |
| $\tau$ | 2.13285 | Expected cell cycle duration (day) | Crivelli et al. [2012] |

The smallest detectable tumour size has been estimated to be roughly 2^30^ ≈ 1 × 10^9^ cells [Carlson, 2003; Schwartz, 1961]. As viral oncology has only been approved for advanced melanoma, we consider tumours with approximately 10^10^ cells. (This corresponds to viral treatment starting 4 tumour doublings after diagnosis.) To ensure that our numerical computations involve numbers of similar magnitude, we measure the number of tumour cells in units of 1010 cells. Given the homeostatic approximation of leukocytes (≈ 6 × 10^9^ cells/L) and roughly 7 litres of blood, we measure the phagocyte concentration in identical units, namely 10^10^ cells.

To illustrate the difference between distributed and discrete delays in the cell cycle duration, we simulate equation (27) without viral therapy for *j* = 6 and the discrete delay case in Figure 2 a). In Figure 2 b), we show the discrete case and the gamma distributed case when *j* = 50. These simulations show that the discrete delay case has a larger basin of attraction than the distributed delay case. This is unsurprising, since for both *j* = 6 and *j* = 50, the result of Corollary 3.4 holds, so all parameter regimes leading to stability of the CFE for the gamma distributed DDE also lead to stability of the CFE in the discrete delay case. Biologically, this corresponds to increased cell cycle duration heterogeneity leading to more robust tumours.

**Figure 2:**
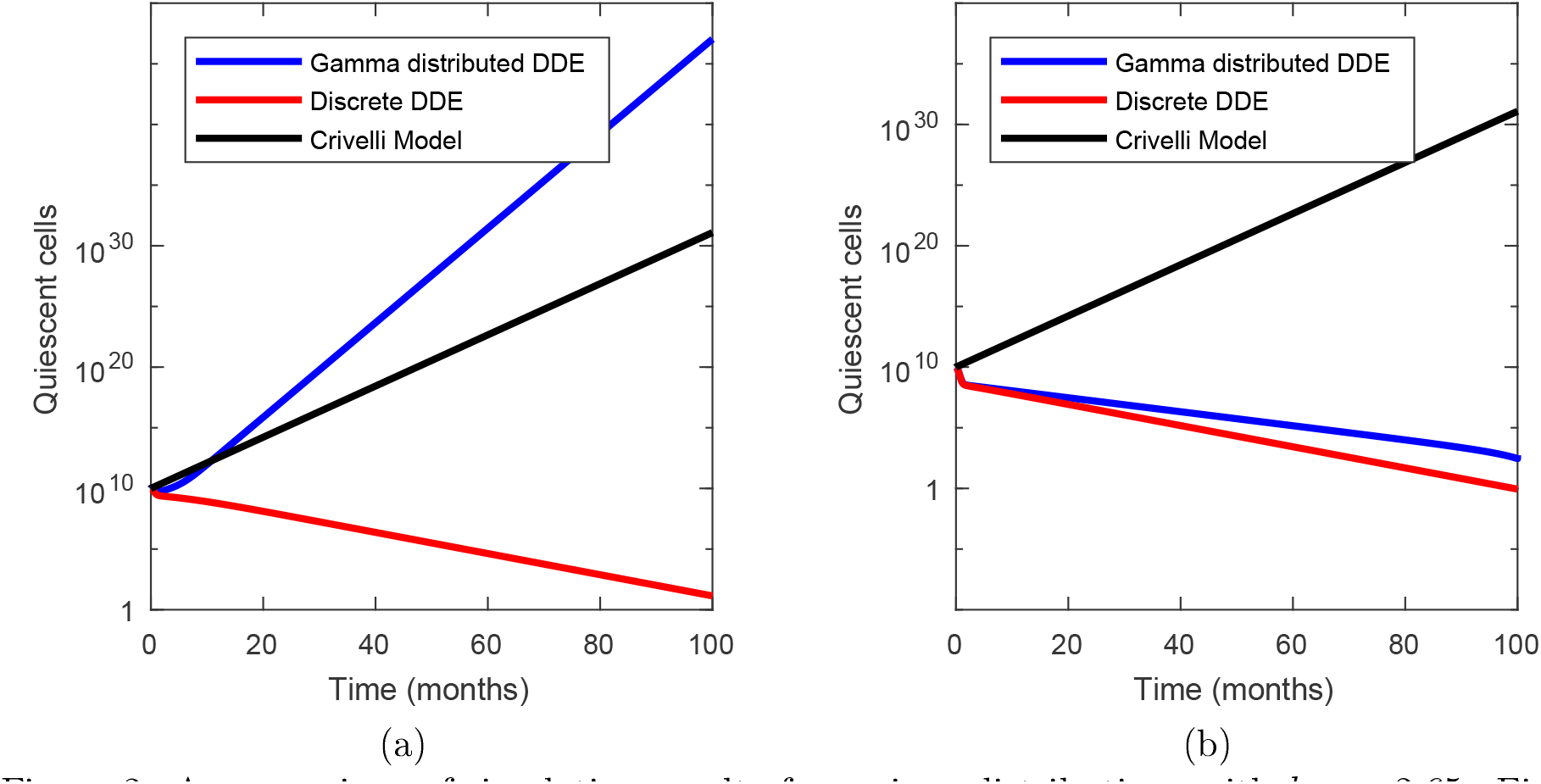
A comparison of simulation results for various distributions with *k_cp_* = 2.65. Figure (a) shows the simulation of equation (1) with a gamma distribution for *j* = 6 in blue, a discrete delay in red and the Crivclli model (equation (37)) in black. Figure (b) shows the simulation of equation (1) with a gamma distribution for *j* = 50 in blue, a discrete delay in red and the Crivelli model (equation (37)) in black.

In Figure 2, we also show the impact of including tumour-immune interaction by comparing our model with that of Crivelli et al. [2012]. We compare the results of our simulation with tumour-immune interaction (*k_p_* = 0.065) with the Crivelli model (*k_p_* = 0) as written in Appendix A. This simulation underlies the importance of tumour-immune interaction in determining disease progression.

In Appendix A, we show that the gamma distribution converges to the degenerate distribution as *j* grows infinitely large, with *τ* > 0 held constant. The case *j* = 1 corresponds to an exponential distribution of cell cycle durations. In what follows, we assume that the distribution of cell cycle durations is neither exponential nor degenerate, so 1 < *j* < ∞. In the numerical simulations that follow, we illustrate a representative case of our results with *j* = 6.

In Figure 3, we simulate the finite dimensional representation of the distributed DDE (1) for different levels of immune recruitment, *k_cp_*, during viral therapy. Figure 3 shows that changing *k_cp_* changes the long-term success or failure of viral treatment. Sufficiently large values of *k_cp_* induce long-lasting remission while smaller values of kcp lead to eventual tumour progression after oncolytic virus treatment.

**Figure 3:**
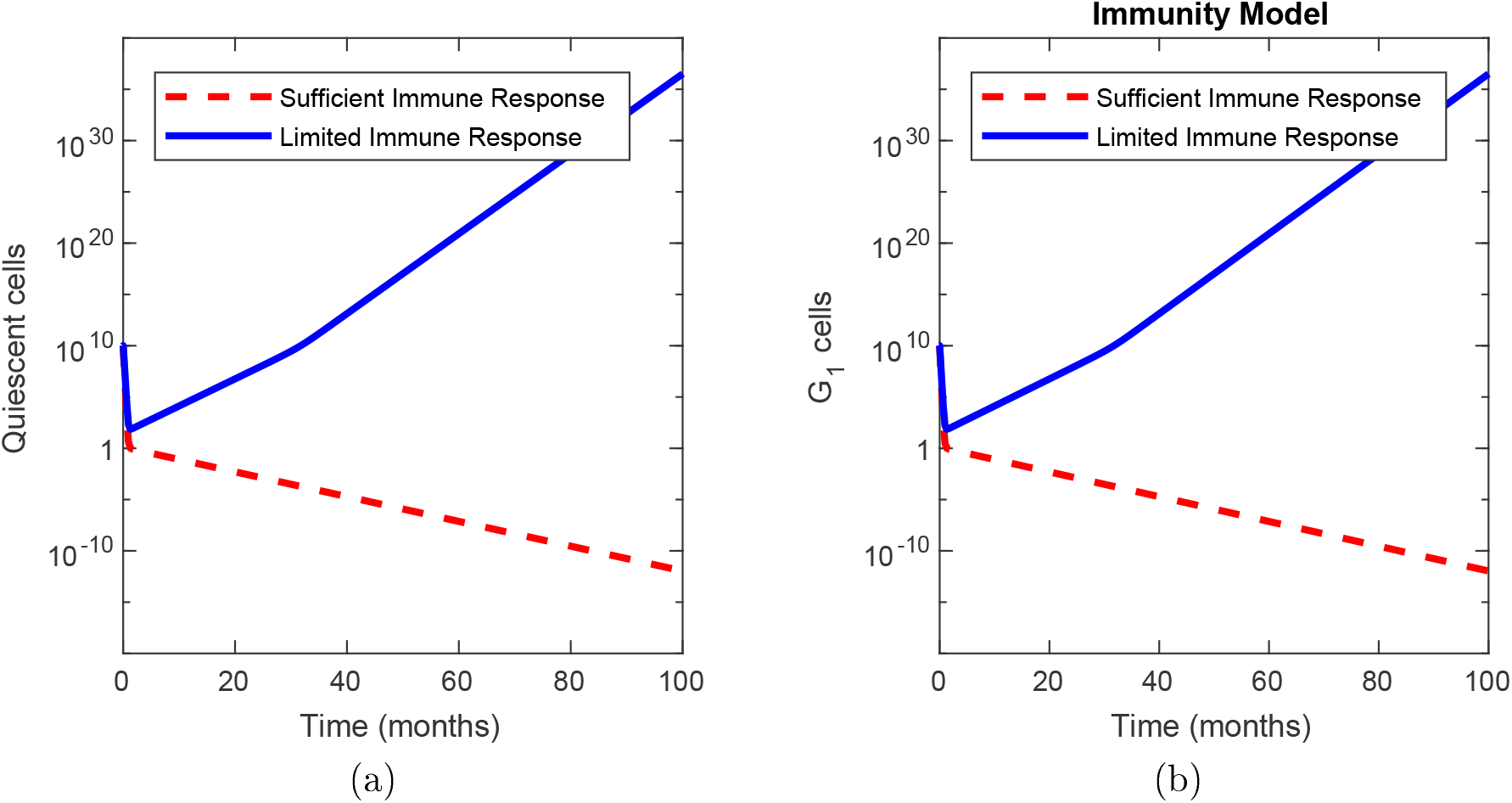
Simulated viral therapy with limited and sufficient immune recruitment. The parameters used in sufficient immune recruitment are given in Table 1. Limited immune recruitment occurs with *k_cp_* = 1.63 and other parameters as given in Table 1.

Figure 4 shows the impact of parameter variability on stability of the CFE. Figure 4 (a) shows that increased immune interaction (*k_p_*) can counteract fast transit between quiescence and mitosis (*a*_1_ and *a*_2_ respectively) to ensure stability of the CFE. Moreover, sufficiently slow entrance into the active phase of the cell cycle (small *a*_2_) also stabilises the CFE. Figure 4 (b) shows that immune recruitment (*k_cp_*) must grow infinitely large to account for less efficient immune-tumour interaction (*k_p_*), while a large death rate during the cell cycle 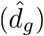 can ensure stability of the CFE regardless of immune involvement. These investigations confirm the impact of immune recruitment and clearance of tumour cells. This result indicates that increasing immune involvement is important in developing therapeutic strategies.

**Figure 4:**
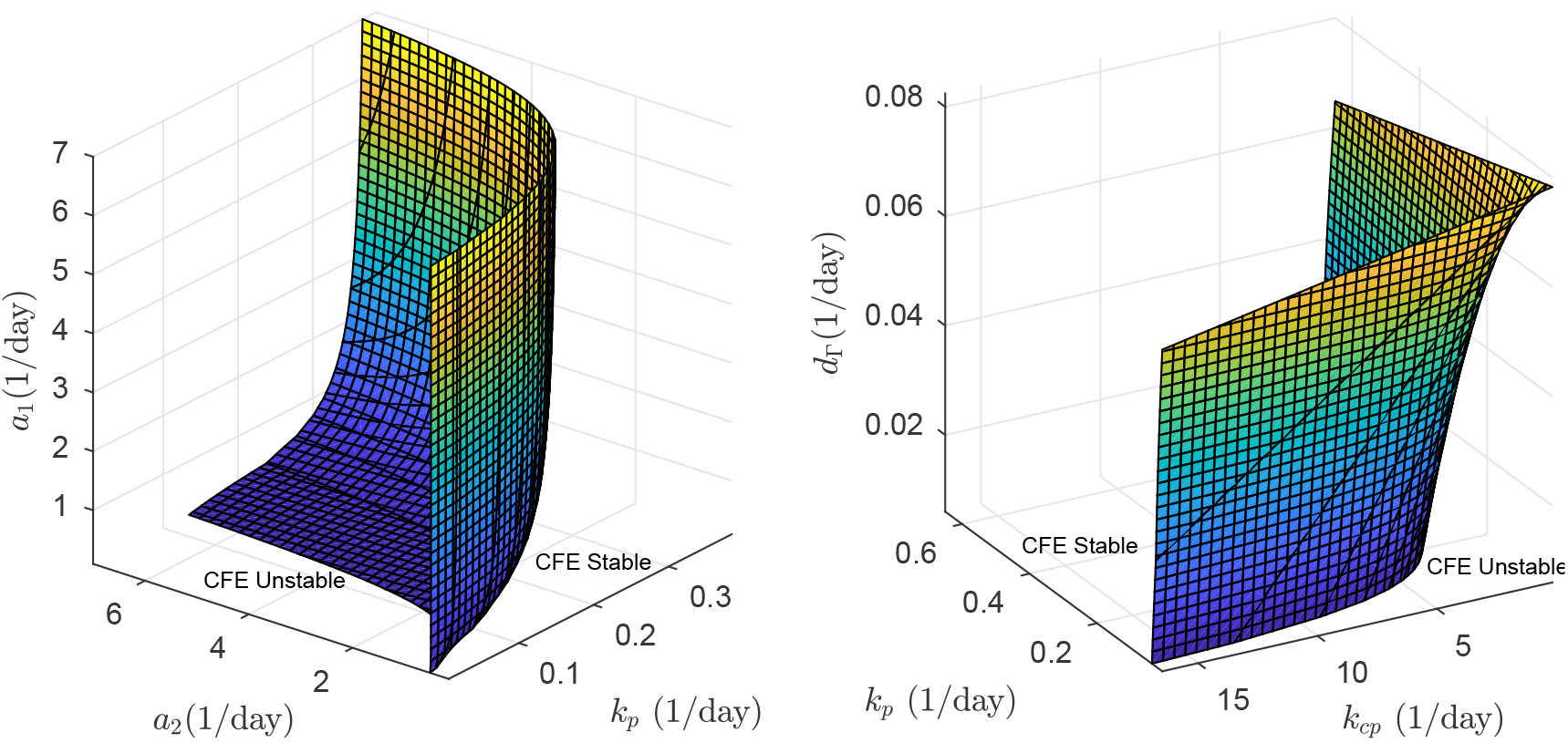
Stability regions for the cancer free equilibrium for various parameter combinations. Figure (a) shows the relationship between the stability of the CFE and the parameters *k_p_, a*_1_ and *a*_2_. Figure (b) shows the relationship between stability of the CFE and the parameters *k_cp_, k_p_* and 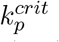.

Finally, Figure 5 shows the relationship between the nonzero equilibrium found in Theorem 3.5 and the parameter *k_p_*. The diagram indicates that the CFE gains stability through a transcritical bifurcation as *k_p_* increases. For 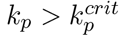 and initial conditions straddling the unstable equilibirum, we see the dependence of asymptotic behaviour on initial conditions. A similar relationship exists between the stability of the CFE and *k_cp_*. Biologically, Figure 5 (b) shows that the same immune system can control small tumours while large established tumours grow unboundedly.

**Figure 5:**
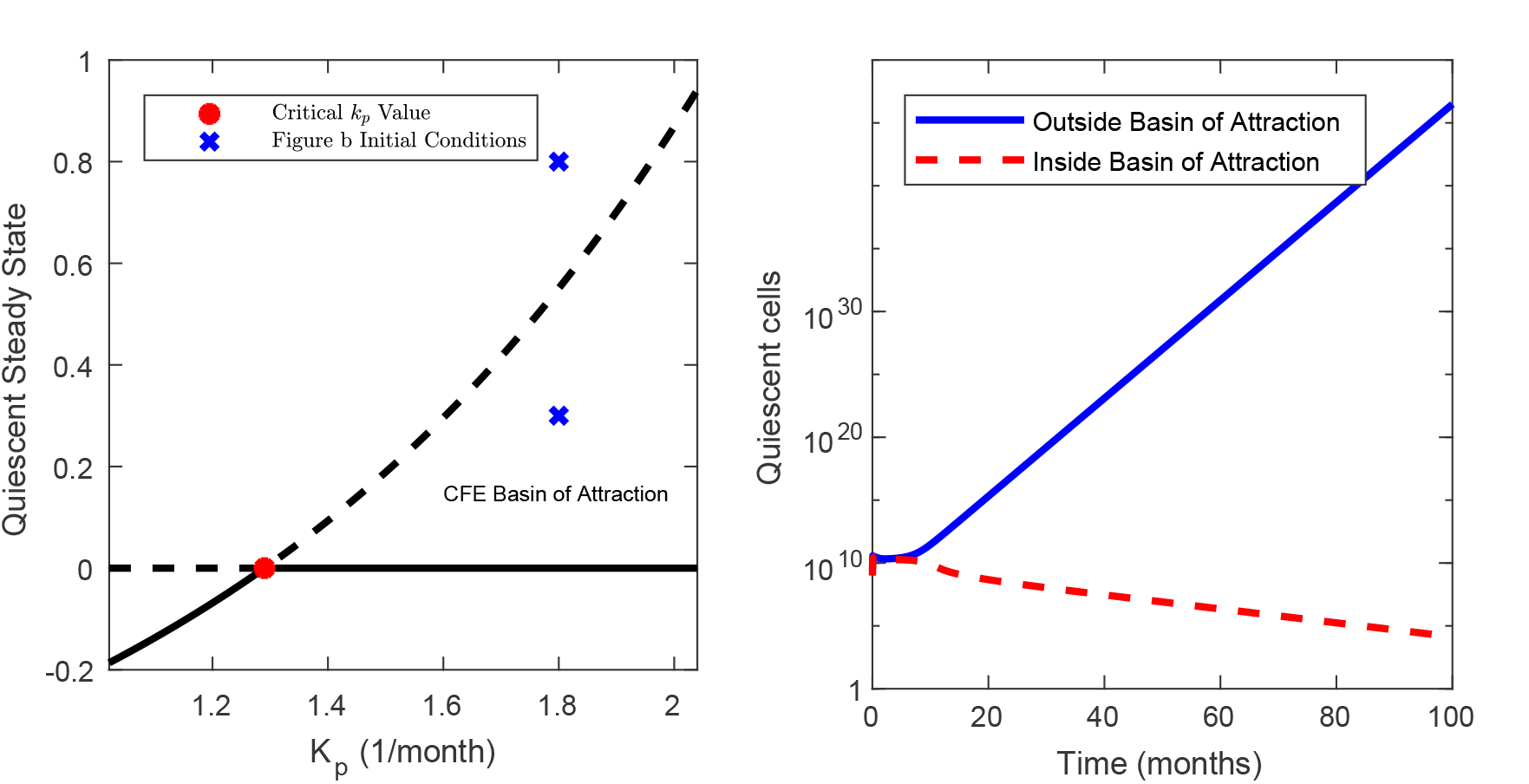
The bifurcation diagram showing a transcritical bifurcation. Figure (a) shows the transcritical bifurcation as *k_p_* increases past 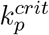 for the quiescent population. The dashed lines represent unstable equilibria and the solid lines denote stable equilibria. Figure (b) the dependence of asymptotic behaviour on initial conditions. The quiescent initial populations used are shown in Figure (a) as crosses.

## 5 Discussion

Malignant tumours are comprised of an extremely heterogeneous population of malignant cells. Oncolytic viruses combat this heterogeneity by exploiting two common characteristics of malignant cells: weakened antiviral immunity and explosive growth rates. Once an oncolytic virus has infiltrated a tumour, lysis of infected cells and immune recruitment combine to eliminate the tumour. Past models of tumour growth and viral oncology have used discrete DDEs to model the cell cycle duration and infection of susceptible cells. However, discrete DDEs enforce a uniform and constant tumour cell cycle time and do not incorporate any aspect of the inherent heterogeneity of malignant cells inside the tumour microenvironment.

In this work, we produced a mathematical model of tumour cell growth that incorporates the heterogeneity of tumour reproduction speed by modelling cell cycle duration as a random variable following a PDF *K*(*t*). This framework is a novel representation of tumour growth and is more physiologically realistic than the discrete delay case. Specifically, variation in tumour cell cycle duration can be seen as a measure of tumour cell heterogeneity. Using linear stability analysis, we established the relationship between the expected number of cells surviving the cell cycle and tumour remission. As we assumed a constant death rate throughout the cell cycle, the expected number of cells surviving the cell cycle is directly related to the distribution of cell cycle durations. The distribution of cell cycle durations and disease progression are explicitly linked in our stability threshold. The stability threshold determines the minimal anti-tumour immune response that ensures that nascent tumours do not persist. This result shows that increasing immune involvement can stabilise the tumour free state regardless of the cancer growth rate.

Our results indicate that lysis of infected cells and increased immune recruitment act synergistically to eliminate tumour cells during viral therapy. Our simulations show that the combination of viral therapy and the resulting immune recruitment function by driving solutions across a separatrix into the basin of attraction of the tumour free equilibrium. If immune recruitment is insufficient to control tumour growth, we predict that viral therapy will drive initial tumour remission that is followed by disease recurrence. Moreover, our results show that viral therapy can act as the external force required to shrink tumours to a size manageable by the immune system, leading to long-term remission. These observations are consistent with clinical results and suggest that oncolytic viruses designed to maximise immune response may have clinical benefits.

Finally, our modelling techniques develop a novel mathematical treatment of tumour cell growth by using a distributed DDE. The distributed DDE considered in this work incorporates the discrete delay case studied by Crivelli et al. [2012] and others for a suitable choice of *K*(*t*) and *k_p_*. In the specific case of a gamma distribution, we derive a novel linear chain technique that incorporates cellular loss throughout the cell cycle. Using this technique, we reduce the infinite dimensional distributed DDE to an equivalent finite dimensional ODE. Our derivation of the equivalent ODE formulation is easily generalisable to physiological processes with exponential growth or decay. The reduction of the distributed DDE to an ODE offers a method whereby models using discrete DDEs can include more physiologically realistic distributed delays without losing the ability to easily simulate the model.

Our modelling framework has certain limitations. The mathematical model greatly simplifies immune recruitment and tumour-immune interactions in favour of an analytically tractable model. The interactions between the legion of cytokines and immune cell types in the tumour micro-environment are not considered in this work, nor have we studied the effect of immune system selection of cancer cells.

This modelling work raises the interesting question of which distribution best models tumour cell cycle durations. Most existing models either use the discrete or gamma distribution to exploit the existing numerical methods to simulate these models. Without data, it is difficult to determine which distribution most accurately models tumour cell cycle durations. Nevertheless, our analytic results are valid for any distribution describing tumour cell cycle durations. In summary, our model incorporates an aspect of tumour cell heterogeneity, makes predictions that are consistent with clinical observations and indicates future avenues of oncolytic virus development.

## Acknowledgements

TC would like to thank the Natural Sciences and Engineering Research Council of Canada (NSERC) for funding through the PGS-D program and the Alberta Government for funding through the Sir James Lougheed award of distinction. ARH is grateful to NSERC, Canada for funding through the Discovery Grant program. Both authors thank Morgan Craig for many helpful discussions.

## Appendix A Reduction to the Crivelli Model and the Discrete Delay Case

We show that the Crivelli model [Crivelli et al., 2012] is a special case of the general distributed DDE model (1) developed in Section 2 without immune recruitment. We do this two ways, first by showing that the discrete DDE model corresponds to the distributed DDE model with a degenerate distribution. Then alternatively by showing that the discrete DDE model can be recovered from the distributed DDE model in a suitable limit when *K*(*t*) is taken to be a Gamma distribution.

Crivelli et al. [2012] do not consider tumour-immune involvement, so we take *k_p_* = 0 in (1). Then, the immune recruitment has no impact on the tumour model, so we drop the differential equations for *P*(*t*) and *C*(*t*). Crivelli et al. [2012] use a discrete DDE to model the cell cycle duration. The simplest way to recover a discrete DDE from a distributed DDE is to let *K*(*t*) = *δ*(*t* − *τ*). Then, equation (7) gives *d_δ_* = *d*_3_. Thus the model (1) becomes 
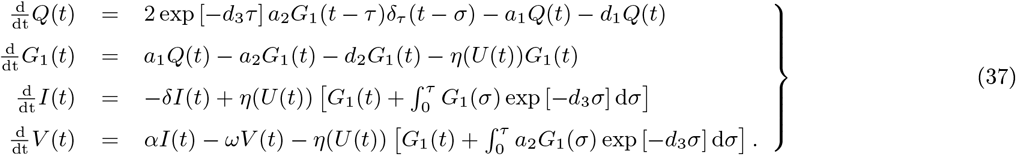

Finally, evaluating equation (2) with *K* (*t*) = *δ*(*t* − *τ*) gives 
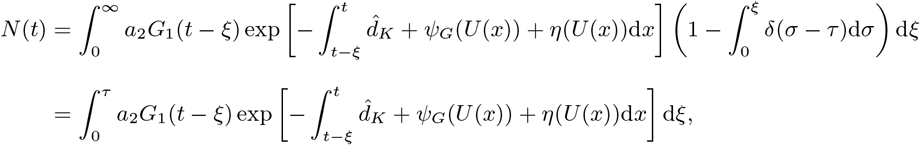
and taking *η*(*U* (*t*)) to be the non-differentiable contact rate 
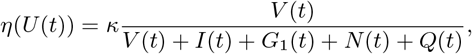
returns the mathematical model in Crivelli et al. [2012]. To illustrate that their results are a special case of ours, we use Theorem 3.3 to determine the stability of the CFE for the Crivelli model. With *k_p_* = 0 and *K*(*t*) = *δ*(*t*−*τ*), it is simple to calculate that Ψ_*G*_1__ = 0 and 
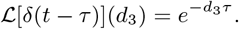

Then the stability condition (22) becomes 
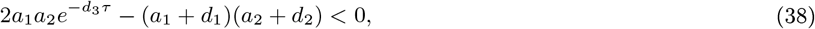
which is exactly the same as found by Crivelli et al. [2012].

We have shown that discrete DDEs can be modelled as degenerate distributed DDEs. Next, we show a distinct method of reducing the general distributed DDE to a discrete DDE by considering a gamma distributed DDE, i.e.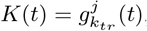, in the limit as *j* → ∞. We parameterise the gamma distribution by choosing *j* ∈ 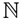 and setting *a_j_ = τ*/*j*. Then, for each integer *j*, the expected duration of the cell cycle is *τ*. Moreover, the standard deviation is given by 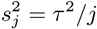 with 
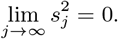

Heuristically, as *j* increases, 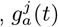 becomes increasingly concentrated about the expected value, *τ*. Formally, the characteristic function of the gamma distribution converges in distribution to the characteristic function of the *δ*(*t* − *τ*) distribution with 
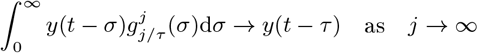
for any test function *y*(*t*). From equation (26), 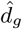 is dependent on the parameter *j* via 
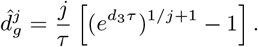

To compute the limit of 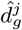 as *j* → ∞, we first note that 
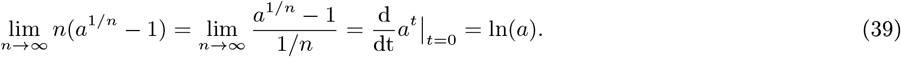

Therefore, 
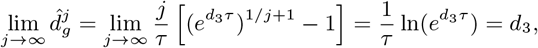
so 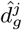 converges to the death rate of the discrete DDE as *j* → ∞.

Finally, we compute the linearisation matrix for the linearised DDE (17) with 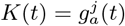: 
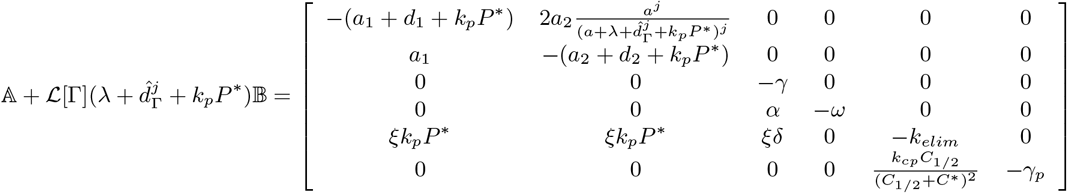
and the corresponding characteristic function, once again using equation (19), 
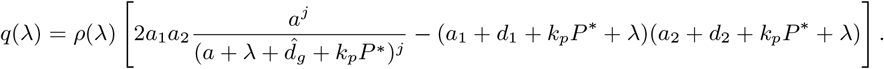

Using Theorem 3.3, the condition for stability of the CFE is 
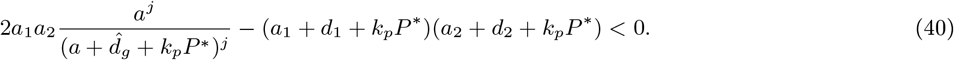

Using *a* = *J*/*τ*, we rearrange this condition to find a cell cycle duration, **τ*_j_*, that ensures local stability of the CFE 
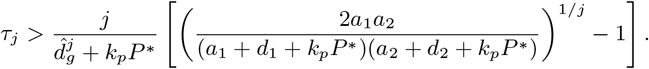

Then, the minimal cell cycle duration for stability, 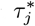, is given by 
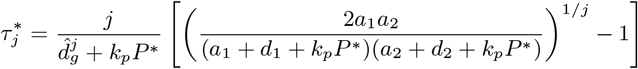
and is dependent on the parameter *j*. Once again, using equation (39), we see that
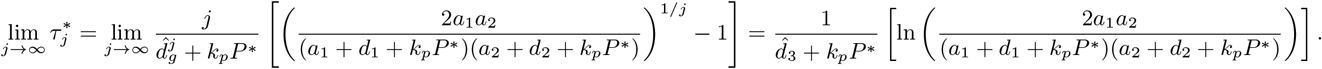

Thus the critical cell cycle duration when 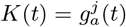 converges to the critical cell cycle duration time in discrete delay case case. Moreover, when 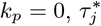 converges to the critical delay time found by Crivelli et al. [2012].

Consequently, the discrete DDE model considered by Crivelli et al. [2012] can be considered a degenerate case of the distributed DDE or as a limit of a gamma type distributions.

## Appendix B Number of Cells in the Cell Cycle

Here, we detail the calculation of the number of cells in the active portion of the cell cycle at time *t*. Fix *ξ* > 0, so the number of cells entering the active portion of the cell cycle at time *t* − *ξ* is *a*_2_*G*_1_(*t* − *ξ*).

Then, at time *t*, the probability that a cell that entered the active portion of the cell cycle at time *t* − *ξ* has not completed the cell cycle is 
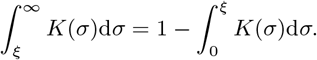

Of the cells that have not exited the active portion of the cell cycle, the fraction that have not died by time *t* is 
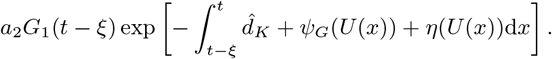

Integrating over all previous times *ξ* gives the total number of cells remaining in the cell cycle. Consequently, the number of cells in the cell cycle at time *t* is 
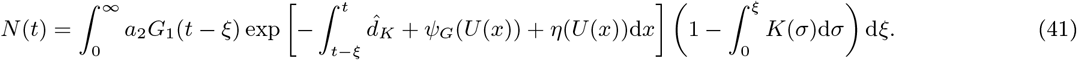

By making the change of variable *v* = *t* − *ξ*, we have the alternative form 
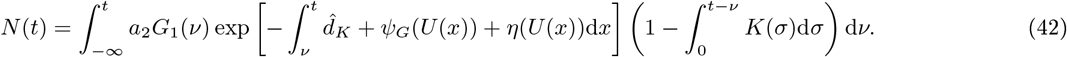

Equation (42) is difficult to evaluate numerically. However, differentiating *N*(*t*) by using the Lebeniz and product rules, we find the distributed DDE for *N*(*t*) 
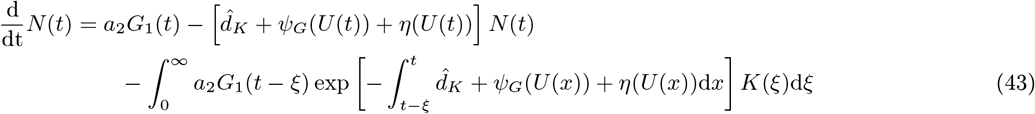
which can be solved numerically. As we have shown in Proposition 4.1, we can replace the distributed DDE (43) with the solution of the transit compartment ODE defined in equation (29) when 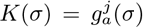. Therefore, in our simulations of equation (27), we calculate *N*(*t*) by solving 
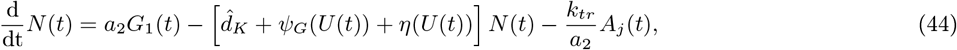
subject to the initial condition from evaluating equation (41) at *t* = 0 by using the lower incomplete gamma function.

